# H3F3A K27M Mutations Drives a Repressive Transcriptome by Modulating Chromatin Accessibility, Independent of H3K27me3 in Diffuse Midline Glioma

**DOI:** 10.1101/2024.05.16.594522

**Authors:** Suraj Bhattarai, Faruck L. Hakkim, Charles A. Day, Florina Grigore, Alyssa Langfald, Igor Entin, Edward H. Hinchcliffe, James P. Robinson

## Abstract

**Background:** Heterozygous histone H3.3K27M mutation is a primary oncogenic driver of Diffuse Midline Glioma (DMG). H3.3K27M inhibits the Polycomb Repressive Complex 2 (PRC2) methyltransferase complex, leading to a global reduction and redistributing of the repressive H3 lysine 27 tri-methylation. This rewiring of the epigenome is thought to promote gliomagenesis.

**Methods:** We established novel, isogenic DMG patient-derived cell lines that have been CRISPR-Cas9 edited to H3.3 WT or H3.3K27M alone and in combination with EZH2 and EZH1 co-deletion, inactivating PRC2 methyltransferase activity of PRC2 and eliminating H3K27me3.

**Results:** RNA-seq and ATAC-seq analysis of these cells revealed that K27M has a novel epigenetic effect that appears entirely independent of its effects on PRC2 function. While the loss of the PRC2 complex led to a systemic induction of gene expression (including HOX gene clusters) and upregulation of biological pathways, K27M led to a balanced gene deregulation but having an overall repressive effect on the biological pathways. Importantly, the genes uniquely deregulated by the K27M mutation, independent of methylation loss, are closely associated with changes in chromatin accessibility, with upregulated genes becoming more accessible. Notably, the PRC2- independent function of K27M appears necessary for tumorigenesis as xenografts of our H3.3K27M/EZH1/2 WT cells developed into tumors, while H3.3/EZH1/2 KO cells did not.

**Conclusion:** We demonstrate that K27M mutation alters chromatin accessibility and uniquely deregulates genes, independent of K27 methylation. We further show the mutation’s role in altering biological pathways and its necessity for tumor development.

**Key Points:**

- We revealed genes regulated by H3.3K27M mutation and PRC2 in DMG.
- H3.3K27M mutation alters chromosome accessibility independent of H3K27me3.
- PRC2-independent effects of K27M mutation are crucial for tumor development.

**Importance of the Study:** This study is the first to demonstrate that H3F3A K27M mutations drive a repressive transcriptome by modulating chromatin accessibility independently of H3K27 trimethylation in Diffuse Midline Glioma (DMG). By isolating the effects of H3.3 K27me3 loss from those of the K27M mutation, we identified common and unique genes and pathways affected by each. We found that genes uniquely deregulated by K27M showed increased chromatin accessibility and upregulated gene expression, unlike other gene subsets affected by PRC2 knockout. Importantly, we determined the PRC2-independent function of K27M is also essential for tumorigenesis, as xenografts of H3.3 K27M/PRC2 WT cell lines formed tumors, while H3.3WT/PRC2 WT and K27M/PRC2 knockout cells did not. This research builds upon and advances prior studies, such as those identifying EZH2 as a therapeutic target in H3.3K27M DMGs, by revealing critical new pathways for gliomagenesis. The translational significance lies in identifying novel therapeutic targets against this aggressive pediatric cancer.

**Graphical Abstract:** 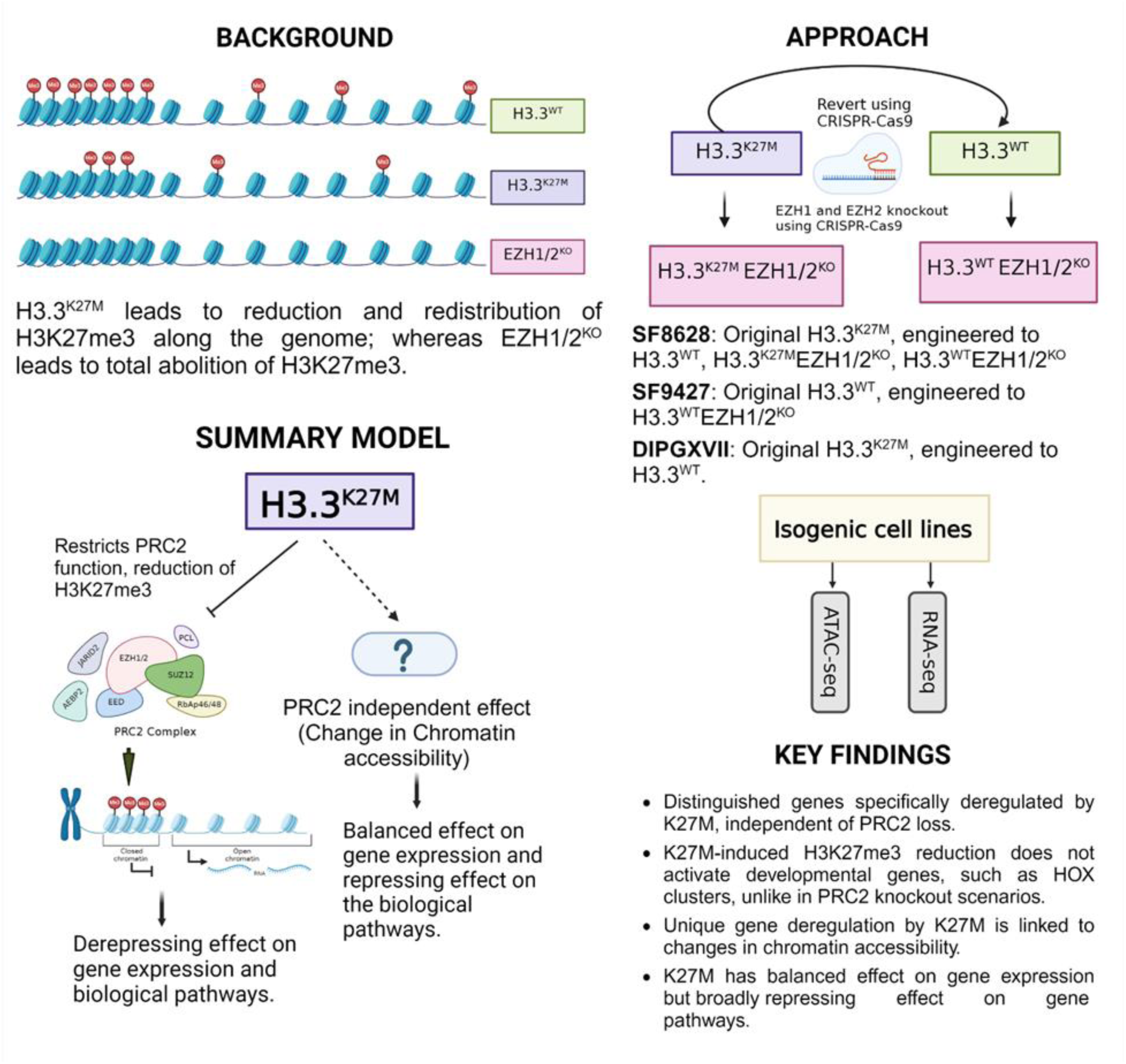

## Introduction

Diffuse Midline Glioma (DMG) poses a significant challenge in pediatric oncology due to its aggressive nature, anatomical location, challenging resection, and resistance to treatment (1). The survival statistics are stark, with median survival post-diagnosis between 9 to 15 months and less than a 2% five-year survival rate (1,2). The identification of heterozygous H3.3 histone A (H3F3A) gene mutations, resulting in the H3.3K27M mutation in about 80% of DMG, marked a critical development in understanding these tumors’ biology (3–8).

The Polycomb Repressive Complex 2 (PRC2) is central to epigenetic gene regulation, primarily through the trimethylation of lysine 27 on histone H3 (H3K27me3) (9–12), which represses genes critical for development, differentiation, and cell proliferation (13–16). This modification is predominantly carried out by the catalytic subunit EZH2, but also by EZH1 which is less efficient than EZH2 (12). Mutations in key members of the PRC2 complex, including enhancer of zeste homolog 2 (EZH2), embryonic ectoderm development (EED), and SUZ12, have been identified across a spectrum of cancers (17,18). Gain-of-function mutations in EZH2, for example, contribute to the progression of cancers such as lymphomas, melanoma, prostate, and breast by promoting the trimethylation of H3K27 and silencing tumor suppressor genes (19–22). Conversely, loss-of-function mutations in EED and SUZ12 lead to a decrease in H3K27me3 levels, which derepresses oncogenes and promotes tumorigenesis in conditions like malignant peripheral nerve sheath tumors (MPNSTs) and certain gliomas (18,23).

The H3.3K27M mutation notably diminishes H3K27me3 levels across the genome, although the exact mechanism is still under investigation (24–33). While the molecular mechanism linking K27M to H3K27me3 reduction is unclear, a recent study has raised questions about just how impactful the loss of H3K27me3 is to glioma formation. Research using doxycycline-inducible embryonic stem cells expressing either wild-type or mutant H3.3K27M and H3.3K27L variants demonstrated that these mutations lead to immature gene expression profiles during cell differentiation (33). Crucially, these changes appeared to be independent of PRC2 function, suggesting that the mutation’s impact on gene regulation and chromatin dynamics involves mechanisms beyond traditional epigenetic silencing pathways. While intriguing, this study was carried out in stem cells only and necessitated follow-up work to establish the presence (and importance) of PRC2-independent, K27M-dependent gene regulation in DMG tumors.

To dissect the dynamics of DMG tumorigenesis, we generated isogenic DMG cell lines with and without the H3.3K27M mutation, employing precise genome editing with CRISPR-HDR. Following this, cells underwent a second round of CRISPR-Cas9 gene editing to knock out EZH1 and EZH2, thus eliminating the H3K27 methylation capacity of the PRC2 complex. This unique set of isogenic cells—H3.3WT/PRC2WT, H3.3K27M/PRC2WT, H3.3WT/PRC2KO, and H3.3K27M/PRC2KO—allowed us to differentiate the epigenetic changes induced by H3K27M from those resulting from altered H3K27me3 levels or PRC2-independent mechanisms. RNA sequencing and ATAC-Seq analyses provided a comprehensive view of gene expression and chromatin accessibility changes, respectively. Importantly, our findings reveal that K27M has a unique influence on transcription resulting from changes in chromatin state, that are entirely independent of PRC2. Loss of functional PRC2 leads to a systemic induction of gene expression and upregulation of biological pathways, even expressing developmentally repressive genes including HOX gene clusters; whereas K27M leads to a balanced gene deregulation; altering a similar number of up/down regulated genes; but having an overall repressive effect on the biological pathways. In vivo, analyses through xenografts demonstrated that loss of K27M prevented tumor growth; however, loss of EZH1/2 also prevented tumor growth in DMG and adult glioma cells. Together, our genomic analysis and animal studies allowed us to elucidate the critical roles of the H3.3K27M mutation and PRC2-mediated methylation in cancer biology and, importantly, suggest that K27M has its own epigenetic function(s) (not mediated through PRC2) that is vital for DMG formation.

## Materials and Methods

### CRISPR/Cas9 editing

To generate SF8628 H3.3 WT revertant cells, a custom Edit-R crRNA (Dharmacon) targeting the K27M region of H3F3A was duplexed with Edit-R tracrRNA (Dharmacon, Cambridge, U.K.) at 94°C for 2 minutes. This complex, along with recombinant Cas9 (Invitrogen, Waltham, MA), H3F3A WT ssDNA, and a GFP-H3F3A WT HDR repair plasmid, was introduced into cells using the Lonza Nucleofector system (Lonza, Basel, Switzerland). Following nucleofection, colonies were screened for GFP expression with an IncuCyte S3 system and verified by Sanger sequencing and allele-specific PCR with agarose gel analysis for heterozygosity.

Guide Sequences:

crRNA: *H3F3A* K27M: 5’- AGAGGGCGCACUCAUGCGAGGUUUUAGAGCUAUGCUGUUUUG-3’

crRNA: *H3F3A* WT: 5’-AGAGGGCGCACUCUUGCGAGGUUUUAGAGCUAUGCUGU UUUG-3’

Homology Directed Repair templates:

ssDNA - *H3F3A* WT: 5’-AGCACCCAGGAAGCAGCTGGCTACAAAAGCAGCTCGCA AGAGTGCGCCCTCTACTGGAGGGGTGAAGAA-3’

H3.3 WT-GFP HDR plasmid: Contains 1210 bp of intron 1, H3F3A cDNA, eGFP cDNA linked by 15 amino acids, a LoxP cassette with IRES and Puromycin resistance, SV40 promoter, and 1195 bp of intron 2.

### Cell lines and culture conditions

The pediatric glioma cell line SF-9427 (H3F3A WT) was obtained from the UCSF Medical Center. The pediatric glioma H3F3A K27M lines, SF-8628 and SF-7761, were acquired from Millipore Sigma (St. Louis, MO). SF-8628, B23, BD12, CH4, and SF-9427, were maintained in DMEM-high glucose (Sigma Cat. No. D6546), 10% FBS, 2 mM L-Glutamine (EMD Millipore Cat. No. TMS-002-C), 1x Amphotericin, and 1x Penicillin-Streptomycin. SF-7761 were grown in Dulbecco’s Modified Eagle Medium DMEM-high glucose, 10% FBS, 2 mM L-Glutamine, 20 ng/mL EGF, 20 ng/mL FGF-2, 1x Amphotericin and 1x Penicillin-Streptomycin. SF-8628, B23, CH4, BD12, SF-9427, SF-7761 were grown at 37°C and 5% CO2.

### EZH1 and EZH2 CRISPR plasmid cloning

EZH1 and EZH2 CRISPR gRNAs were designed using GUIDES software (http://guides.sanjanalab.org/) (45). gRNAs were cloned into the BsmBI digested linear Lenti-

Cas9-gRNA-GFP vector (Addgene #124770) as described earlier (Giuliano et al. 2019). Plasmids were propagated by transforming into Stbl3 E. coli.

CRISPR gRNA sequences

EZH1 gRNA 1: TTGGTAGTTGTACACTTGTG

EZH2 gRNA 1: TAGCAAAGATGCCTATCCTG

EZH2 gRNA 2: CAGGATGAAGCAGACAGAAG

### Lentivirus packaging, transduction, and selection of knockout clones

HEK293T cells were transfected with Lenti-Cas9-gRNA-GFP along with packaging plasmids (pMD.2 and PAX) using the calcium phosphate method (46). 24 hrs of post transfection media was changed and virus containing supernatant was collected at 48 to 72 hours post-transfection. Lentivirus were filtered through a 0.45 μm syringe and transduced the target cells with 4-10 μg/mL polybrene. Culture media was changed after 24 hrs of transduction. GFP expressed target cells were sorted and single cell colonies were picked and expanded. The knockout clones were confirmed by western blot and sanger sequencing.

### Western Blotting

Western blotting was performed as described previously (47). See supplementary materials for list of antibodies.

### RNA-seq analysis

Total RNA was extracted using Qiagen’s RNeasy kit and sent to Active motif for 150 bp paired end sequencing by Illumina NovaSeq platform. The analysis of RNA-seq data was conducted using a detailed in-house bioinformatics pipeline. Briefly, the quality of the raw sequencing data was initially assessed by FastQC (version 0.12.1) (48). Despite the common practice of bypassing trimming in favor of the STAR aligner’s soft-clipping capabilities, we found that trimming adapters and poor-quality bases with fastp (version 0.23.2) (49) before alignment improved the unique read mapping percentage. Sequencing reads were aligned to the human reference genome (GRCh38.p14) using STAR (version 2.7.11a) (50). The Rsubread package (version 2.16.0), specifically through the featureCounts function (51), was then employed to assign aligned reads to genomic features, converting BAM format alignment files into raw gene read counts. Differential gene expression analysis was performed utilizing DESeq2 (version 1.40.2) (52) to identify genes that were significantly upregulated or downregulated between mutant cells and their isogenic wild-type cells, defining significance by a log 2 fold change exceeding 1.5 and an adjusted p-value below 0.05.

### Pathway/network analysis/venn diagram

Enriched biological pathways within the differentially expressed genes were determined using gene ontology (GO) analysis in the clusterProfiler package (53). Data visualization was enhanced through the use of volcano plots with EnhancedVolcano (54) and Venn diagrams via the VennDiagram package (55). To further evaluate the impact of H3.3K27M mutation on biological pathways, the clusterProfiler package was utilized for gene set enrichment analysis (GSEA) using all the C2 curated gene sets from the Molecular Signature Database (MSigDB). Gene networks for up and down regulated genes were built using STRINGdb package (56). Network analyses were conducted with a confidence score threshold set at 400 to balance the inclusivity of protein-protein interactions (PPIs) against the risk of false positives.

### ATAC-seq analysis

The ATAC-seq analysis was initiated with paired-end 42 (PE42) sequencing reads, obtained through Illumina NovaSeq platform by Active Motif. Sequencing reads were aligned to the human reference genome (GRCh38.p14) using Burrows-Wheeler Aligner (BWA mem) algorithm under its default parameters (57). For further analysis, mapped reads were filtered through Illumina’s purity filter, retaining only the reads with no more than two mismatches, and mapped uniquely to the genome, while removing the PRC duplicates. To identify peaks indicative of open chromatin regions, MACS3 peak calling algorithm was employed (58).

## Results

### Establishment of Isogenic H3.3K27M and H3.3 WT DMG Cell Lines

Using CRISPR/Cas9 editing targeting the H3.3K27M region of the H3F3A gene in a DMG patient-derived cell line (SF8628), we generated SF8628 H3.3 wild-type (WT) revertant cells. Sanger sequencing confirmed heterozygosity in GFP-positive clones. Edited clones were also validated with immunoblotting and immunofluorescence, which confirmed the WT reversion of K27M mutants and restoration of H3K27me3 (Figure 1A and Supplementary Figure 1). Immunoblotting and antibody-based protocols may not be reliable due to the presence of neighboring post-translational modifications (PTMs) that can make it difficult to distinguish and quantify modification states (34–36). So, we further validated our gene editing by mass spectrometry to measure the relative abundance of the most common histone PTMs in H3.3K27 mutant DMG cell lines and a CRISPR revertant control.

**Figure 1.**
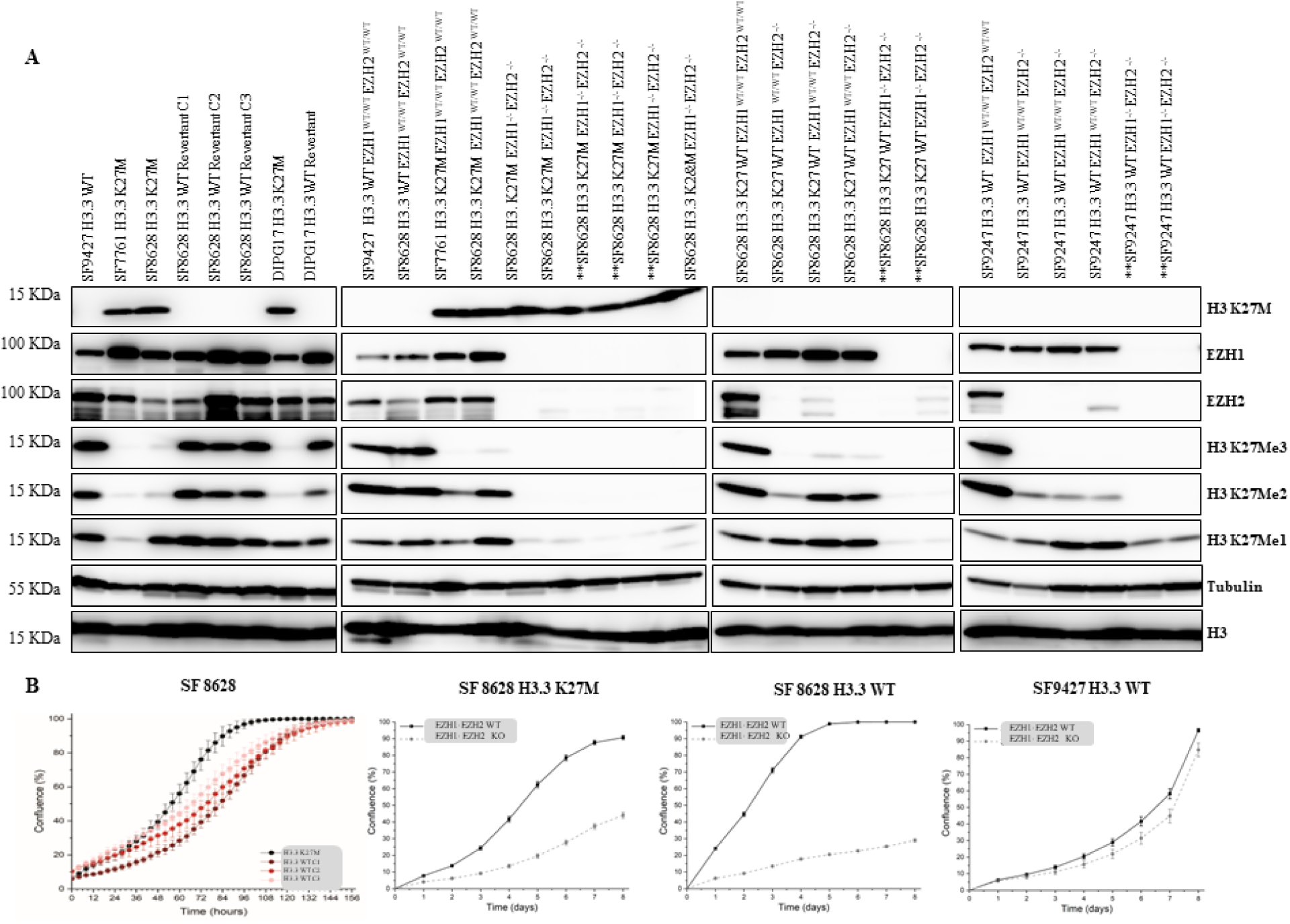
Genetic Manipulations and Phenotypic Effects. (a) Western Blot Analysis Showing CRISPR-Cas9 HDR-mediated Reversion of H3.3K27M Mutation to Wildtype; Knockout of EZH1 and EZH2 Reduces Methylation at H3K27. (a) Growth Rate Analysis of H3.3 Wildtype Revertant Clones Demonstrating Slower Growth. Impact of EZH1 and EZH2 Knockout on Growth Rate in K27M and Wildtype Revertant Cells, Showing Reduced Growth in K27M and Revertant Cells, but Not in Wildtype H3.3 Cells.

Our analysis identified significant differences in histone PTMs between SF8628 K27M mutant DMG cells and CRISPR revertant controls. Reduced methylation at lysine 27 was observed on both H3.1 and H3.3 histone variants in K27M cells (me1: −28.1%, me2: −66.1%, me3: −79.8%, Supplementary Table 1, Supplementary Figure 1) compared to H3.3 WT revertants. Our comparative data from two additional DMG-derived cell lines (the H3.3 WT SF9427 cells, and the H3.3K27M SF7761 cells) emphasizes the pronounced impact of the K27M mutation on histone methylation. We did not observe any significant changes in H3 K27 acetylation, contrast to a trend of increased acetylation reported by other studies (30,31). Interestingly, we found that the H3.3K27M mutant constituted 50% of the H3.3 protein in SF7761 cells, suggesting either upregulation or copy number variations.

Lastly, we conducted a growth analysis of all CRISPR clones using continuous monitoring with the IncuCyte S3 in-incubator cell imaging system. Our findings revealed that H3.3 WT revertant cells exhibited a slight but significant reduction in growth compared to parental cells (Figure 1B).

### Establishment of Isogenic EZH1^-/-^ and EZH2^-/-^ DMG Cell Lines

Using CRISPR/Cas9 editing, we generated EZH1-/- and EZH2-/- knockout SF8628 cells harboring the H3.3K27M mutation, SF8628 cells with wild-type H3.3 (3 pooled clones), and SF9247 cells. After single-cell selection, gene-edited clones were confirmed through immunoblotting, Sanger sequencing, and immunofluorescence (see Figure 1A and Supplementary Figure 2). Our results showed that the co-deletion of EZH1 and EZH2 resulted in the elimination of H3 K27me2 and me3. These findings are consistent with those previously reported in EZH1-/- and EZH2-/- cells (10,37). Again, we utilized an IncuCyte S3 to measure cell growth and proliferation of all CRISPR clones. EZH1-/- and EZH2-/- cells exhibited a significant reduction in growth in SF8628 WT and K27M mutant cells but not H3.3 WT SF9247 cells (Figure 1B).

### Analysis of gene expression changes induced by the H3.3K27M mutation and EZH1/EZH2 knockout

RNA and DNA were extracted from the isogenic H3.3 mutant and revertant, EZH1/2 wild type and knockout cell lines as described and characterized above, as well as from paired isogenic DIPGXVII H3.3K27M mutant and wild type revertant cells (kindly gifted by Paul Knoepfler (38)). These samples were analyzed by RNA sequencing and ATAC sequencing. The PCA plot using RNA-seq data (Figure 2A) demonstrates distinct gene expression profiles across various cell types and mutations evident by the clusters based on cell types, and mutation status. A series of volcano plots visually summarizes the snapshot of transcriptional changes in H3.3K27M and EZH1/2 knockout (KO) mutants compared to their isogenic wild-type counterparts. These plots reveal distinct patterns of gene expression alterations between these mutations. In the volcano plots for the EZH1/2 KO mutants (Figure 2B,D,F), there is a notable predominance of genes on the right side of the vertical threshold line, indicating a significant upregulation compared to the wild-type. This pattern suggests an induction of gene expression following the loss of the EZH1/2 components of the PRC2 complex, which typically acts to repress gene transcription. Conversely, the volcano plots for the K27M mutants (Figure 2C,E) display a more balanced distribution of upregulated and downregulated genes. These observations provide preliminary evidence that while the K27M mutation results in balanced deregulation of gene expression, the knockout of EZH1/2 components distinctly leads to gene induction.

**Figure 2.**
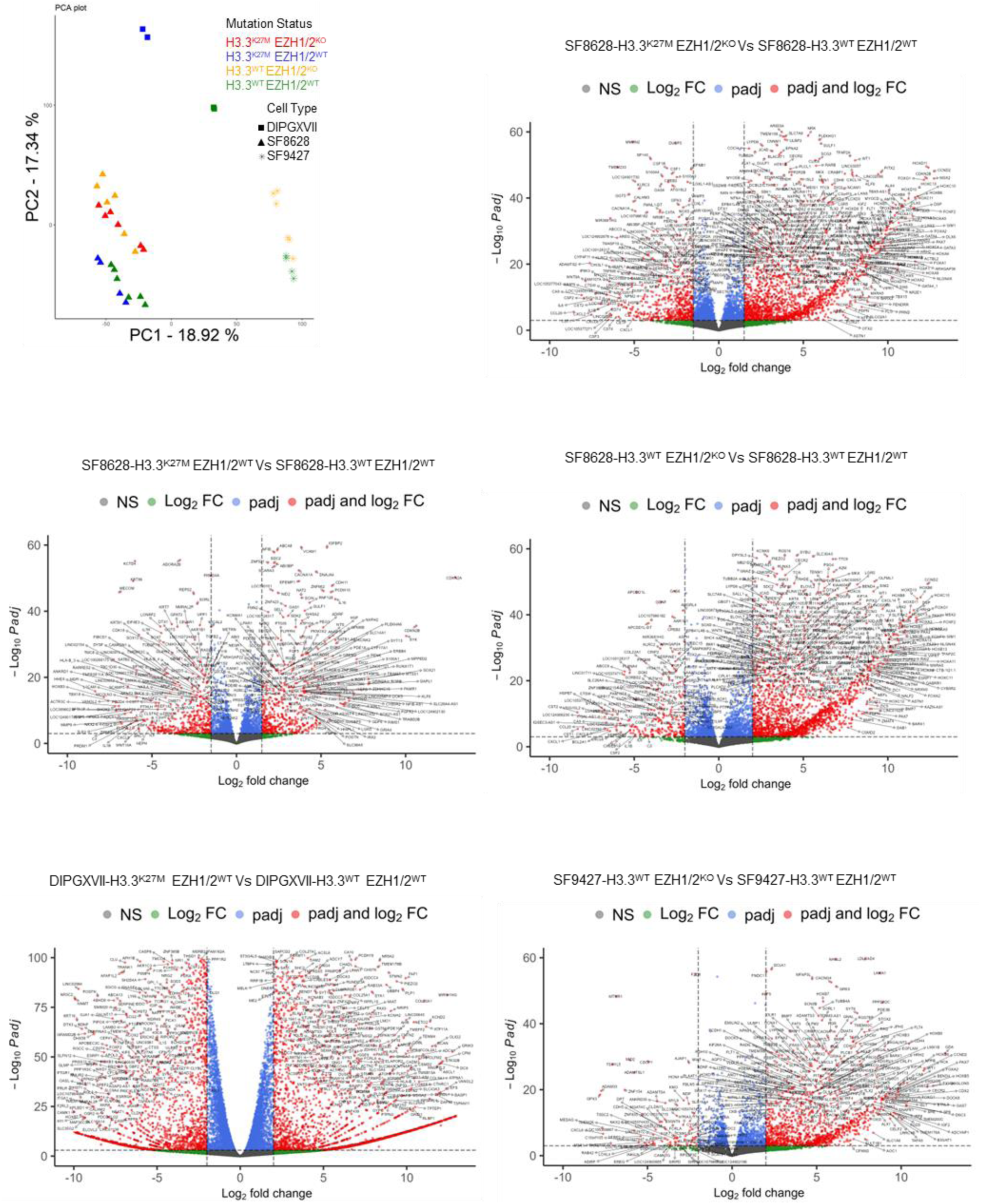
Differential Gene Expression in H3.3K27M Mutation and EZH1/2 Knockout vs. Isogenic Wild Type. A) PCA plot demonstrates that the samples cluster based on cell type and mutation status. B-F) Volcano plots illustrating RNA-seq results comparing K27M and EZH1/2 KO mutants to their respective isogenic wild-type controls. Log2 fold change (FC) is plotted on the x-axis to show expression changes, and negative log10 of the adjusted p-value (padj) on the y- axis for statistical significance. Red points signify genes with statistically significant changes (padj < 0.05) and a log2 FC beyond ±1.5. Vertical dashed lines mark the threshold of log2 FC at ±1.5; red points to the left indicate downregulated genes, and to the right are upregulated genes in mutants relative to wild-type controls. Gene names are annotated to minimize overlap (max.overlaps set to 50), facilitating readability of significant gene markers.

### H3.3K27M mutations cause an overall repression of biological pathways, while their effect on gene expression is neutral

Our transcriptomic analysis revealed distinctive patterns of gene regulation in response to the K27M mutation and EZH1/2 knockouts. In SF8628 cells exhibiting K27M mutation, the ratio of up- and down-regulated genes is 0.99, indicating that a similar number of genes are up- and down-regulated in response to the K27M mutation (Supplementary Table 2). In contrast, the effect of EZH1/2 knockout is derepressing since the number of upregulated genes is 1.4 and 2.5 times higher than downregulated genes in K27M-EZH1/2KO and K27WT-EZH1/2KO cells respectively. These findings are consistent and validated in other cell models: SF9427-EZH1/2KO and DIPGXVII-K27M cells when contrasted with their isogenic wild types (Supplementary Table 2).

We further differentiated genes that are dysregulated by K27M mutation and not by the loss of PRC2 complex. These genes are crucial in terms of DMG biology, as over 80% of DMG cases have this mutation and the most aggressive forms of DMGs almost always bear this driver mutation K27M. Using the Venn diagrams in (Figure 3A,B), we identified a subset of genes uniquely dysregulated by the K27M mutation, distinct from those affected by EZH1/2 knockout. We then performed pathway enrichment analysis on these K27M- specific genes. Notably, the dysregulated genes uniquely associated with the K27M mutation are predominantly contributing to pathways that are downregulated (Figure 3C). This implies that the K27M mutation leads to an overall repression of biological pathways even though it led to a similar number of up- and down-regulated genes. This is further substantiated by similar trends observed in DIPGXVII-K27M cells. The Gene Set Enrichment Analysis (GSEA) in DIPGXVII cells mirrors this downregulation, providing an additional layer of validation (Figure 3J).

**Figure 3.**
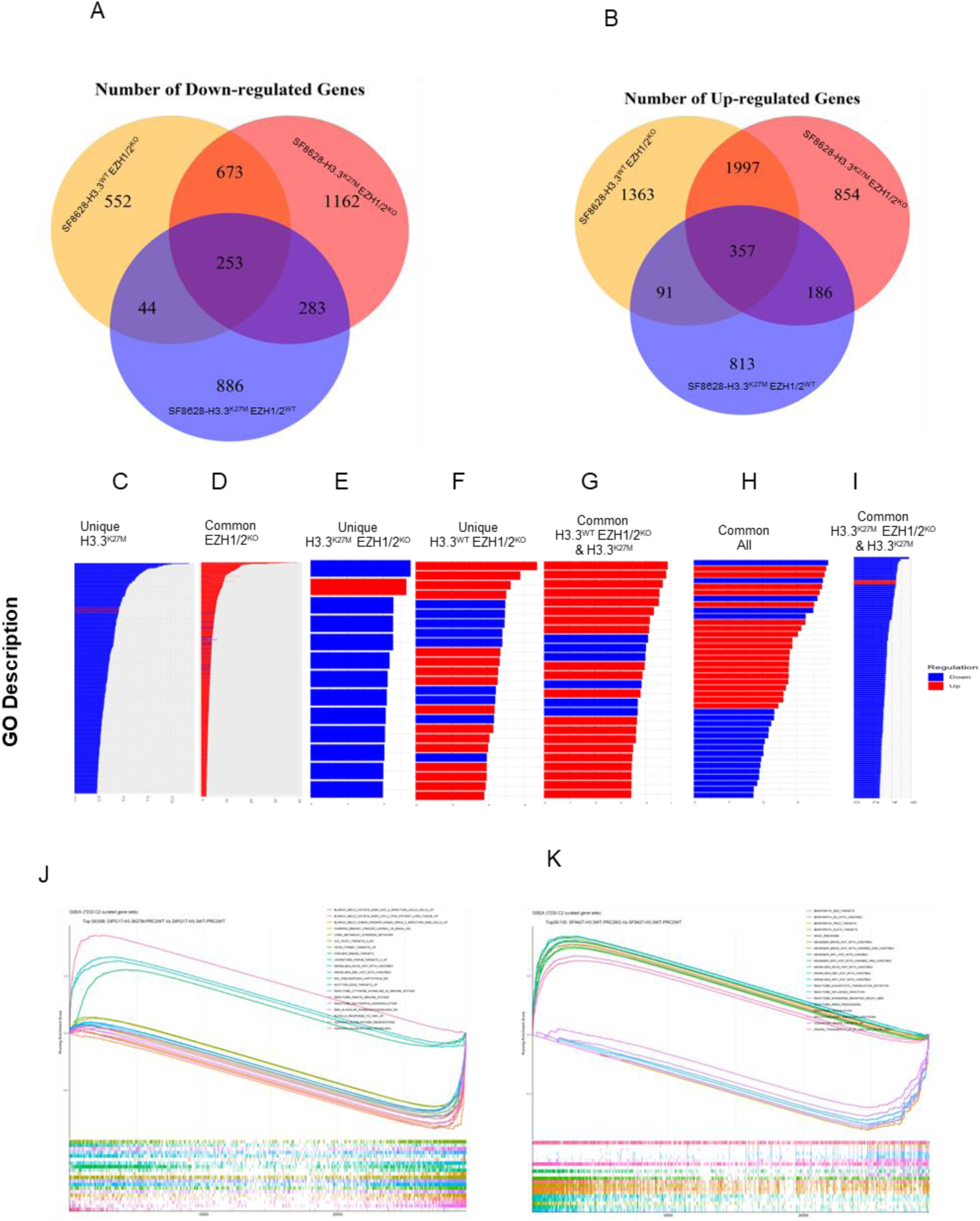
H3.3K27M Mutations Cause an Overall Repression of Biological Pathways while the Effect on Gene Expression is Neutral. Differential gene expression and pathway responses in K27M and PRC2 mutants. (A) Venn diagram illustrating unique and shared down-regulated and (B) up-regulated genes in SF8628 cells with K27M mutation, and EZH1/2KO (PRC2 loss). Gene ontology pathway enrichment analysis for different set of genes unique or common to different mutants from the above venn. Upregulated pathways are shaded in red, and downregulated pathways are shaded in blue. Gene Set Enrichment Analysis (GSEA) plot using 7233 C2 curated gene sets from MolSigDB for (e) DIPGXVII cells highlighting pathways associated with the K27M mutation; and (f) SF9427 cells with EZH1/2 knockout.

Conversely, genes commonly dysregulated by EZH1/2 knockout, regardless of the K27M mutation, predominantly drive pathway upregulation (Figure 3D). This finding aligns with the known repressive role of the PRC2 complex in gene expression and is evident in the pathway activation observed in SF9427-EZH1/2KO cells (Figure 3K). Thus, the loss of functional PRC2 appears to uniformly alleviate repression across the transcriptome. The list of significant GO terms from different overlapping categories is provided in the supplementary table 3.

### K27M distinctly alters chromatin landscape independent of methylation loss

To understand whether the alteration of gene expression we observed in K27M and EZH1/2 knockout cells can be explained by the change in chromosome accessibility, we conducted an analysis of our ATAC-seq datasets. Initial analyses, illustrated by scatter plots, revealed a modest yet consistent correlation between RNA-seq and ATAC-seq log2 fold changes throughout the genome (Figure 4A-C). Specifically, R-squared values of 0.33, 0.39, and 0.47 were recorded for K27M, K27M- EZH1/2, and WT- EZH1/2 cells compared to wild type, respectively, suggesting a moderate association between chromatin accessibility alterations and gene expression changes induced by these mutations.

**Figure 4.**
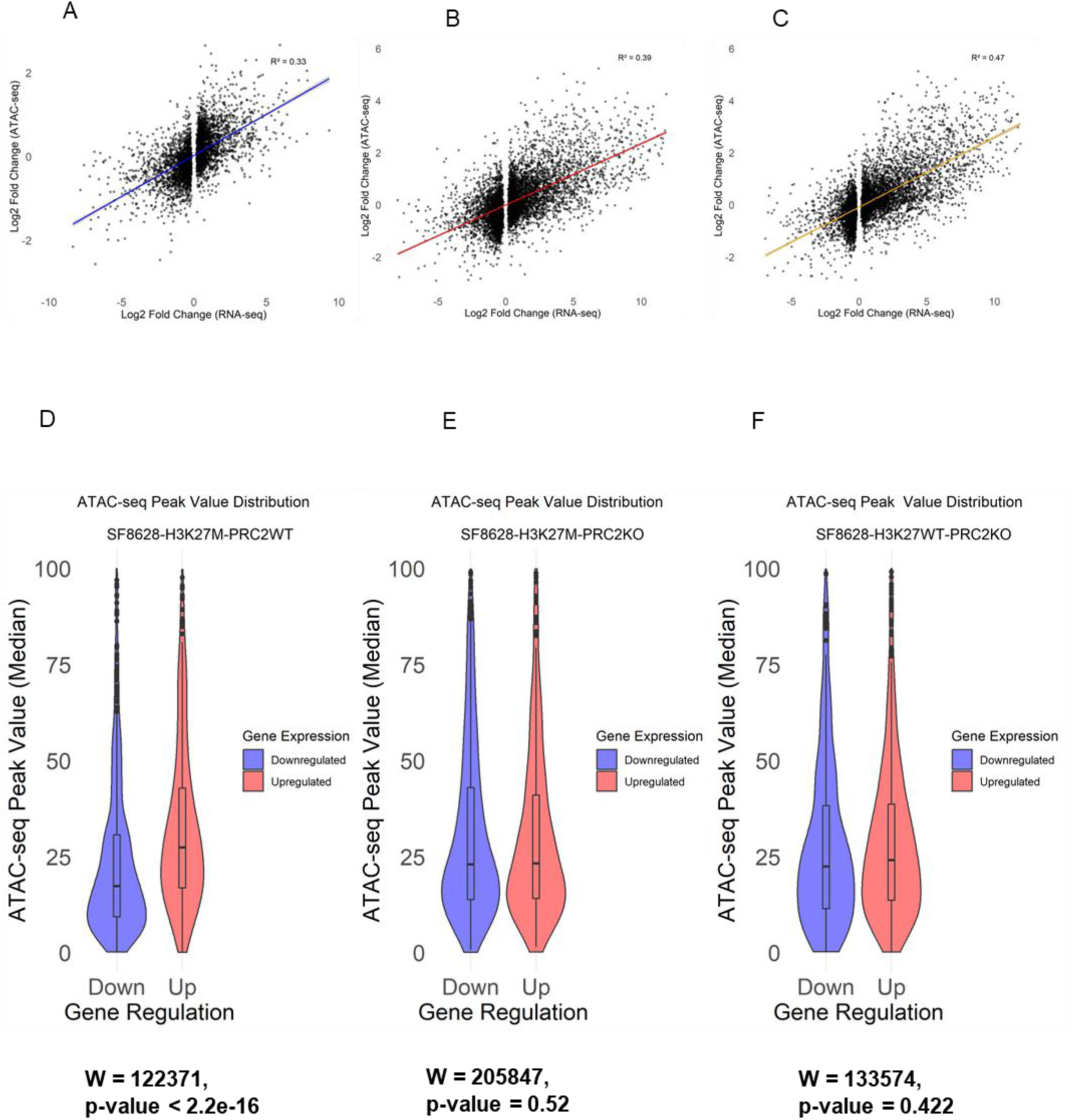
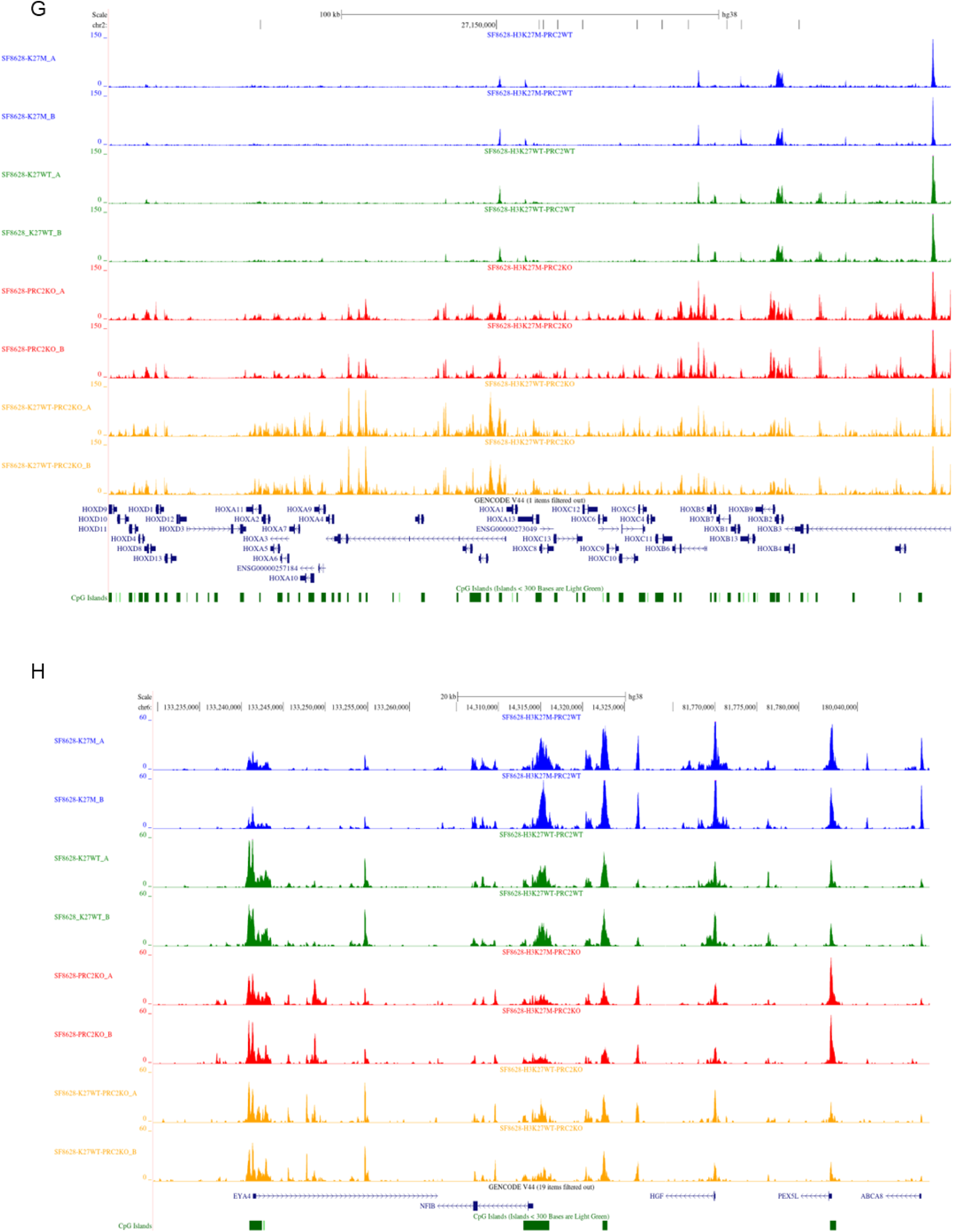
K27M alters chromosome accessibility independent of methylation. Integrative analysis of gene expression and chromatin accessibility across SF8628 mutant cells. a-c Scatter plot demonstrating the correlation between log2 fold changes in RNA-seq and ATAC-seq data for SF8628 cells with different mutation types (A) SF8628-H3K27M-PRC2WT (B) SF8628-H3K27M-PRC2KO and (C) SF8628-H3K27WT-PRC2KO; compared to the wild-type. Each point represents an individual gene. The colored lines indicates the linear regression fit, and the R² value denotes the proportion of variance in ATAC-seq data explained by RNA-seq data. d-f Violin plots showing the distribution of median ATAC-seq peak value for genes uniquely dysregulated in (D) SF8628-H3K27M-PRC2WT (E) SF8628-H3K27M-PRC2KO (F) SF8628-H3K27WT-PRC2KO cells. Down-regulated genes are represented in blue, and up-regulated genes are shown in red. (G-H) UCSC browser snapshots of ATAC-seq peak values for selected genes including (G) HOX gene clusters, which are upregulated in PRC2 knock outs; and (H) NFIB, HGF, PEX5L, and ABCA8, which are up-regulated in K27M. K27M, K27WT, K27M-PRC2KO, and WT-PRC2KO are represented in blue, green, red, and orange color respectively.

Focusing on the impact of the K27M mutation, our research provides compelling evidence of its direct influence on chromatin accessibility. The violin plots demonstrate a significant divergence in ATAC-seq peak values between upregulated and downregulated genes (Figure 4D), supporting the hypothesis that the K27M mutation affects gene regulation by altering the chromatin landscape. The Wilcoxon rank-sum test, chosen for the non-normal distribution of the data, reveals a moderate yet significant effect size, indicating that genes upregulated due to the K27M mutation coincide with increased chromatin accessibility.

In contrast, the PRC2 knockout shows a different pattern. Genes uniquely dysregulated due to the loss of EZH1/2 do not show a significant shift in chromatin accessibility between the up and down-regulated genes (Figure 4E,F), suggesting that gene regulation changes in these cells are not directly caused by changes in the chromatin state. This finding suggests that EZH1/2 knockout has a widespread impact on gene expression that is independent of chromatin accessibility alterations.

Further, we observed a pronounced differential impact of EZH1/2 knockout compared to K27M mutation on the chromatin state of HOX gene clusters. PRC2 disruption led to a substantial increase in open chromatin regions across these loci (Figure 4G), which was paralleled by a significant upregulation of HOX gene expression (Figure 2B,D,F). In contrast, the K27M mutation did not exhibit a similar capability to induce chromatin relaxation within these genes, a finding that was mirrored in their relatively stable expression levels as per our RNA-seq data. The H3K27M mutation exerts a dominant-negative effect on PRC2 function, inhibiting its enzymatic activity. However, this doesn’t lead to the complete loss of H3K27me3 across the genome. Rather, it results in a global reduction and redistribution of H3K27me3, which might selectively affect gene expression. And it appears that HOX genes (which are crucial regulators of embryonic development and cell differentiation) remain comparatively transcriptionally silent in the presence of the K27M mutation (Figure 4G). These genes, typically marked by H3K27me3 for repression during development, retain their silenced state, indicating that the reduction of the repressive marks brought about by the K27M mutation is not sufficient to activate their transcription.

Intriguingly, this trend of chromatin and expression modulation by K27M mutation, as opposed to PRC2 knockout, was not universally applied. For instance, genes such as NFIB, HGF, PEX5L, and ABCA8, which are upregulated and thus demonstrate more open chromatin in the presence of K27M mutation, did not follow the same pattern under EZH1/2 knockout (Figure 4H). This emphasizes the complexity of K27M’s epigenetic regulation, suggesting a selective rather than a broad, indiscriminate influence on chromatin states.

### Summary of distinct GO enriched pathways and gene networks associated with K27M versus EZH1/2 KO

The Gene Ontology (GO) enrichment analysis illustrates a stark contrast between the biological processes influenced by genes uniquely downregulated versus those upregulated in K27M mutants. For the downregulated genes, a diverse array of developmental processes emerges, spanning embryonic organ development, neural growth, and muscle tissue development (Figure 5A). In stark contrast, the upregulated genes are associated with a notably narrower spectrum of enriched pathways, all linked to the extracellular matrix (Figure 5B). The specificity of these pathways, particularly the organization of extracellular structures, might reflect a targeted deregulation by K27M mutation, which could facilitate altered cell-matrix interactions contributing to the invasive properties of gliomas.

**Figure 5.**
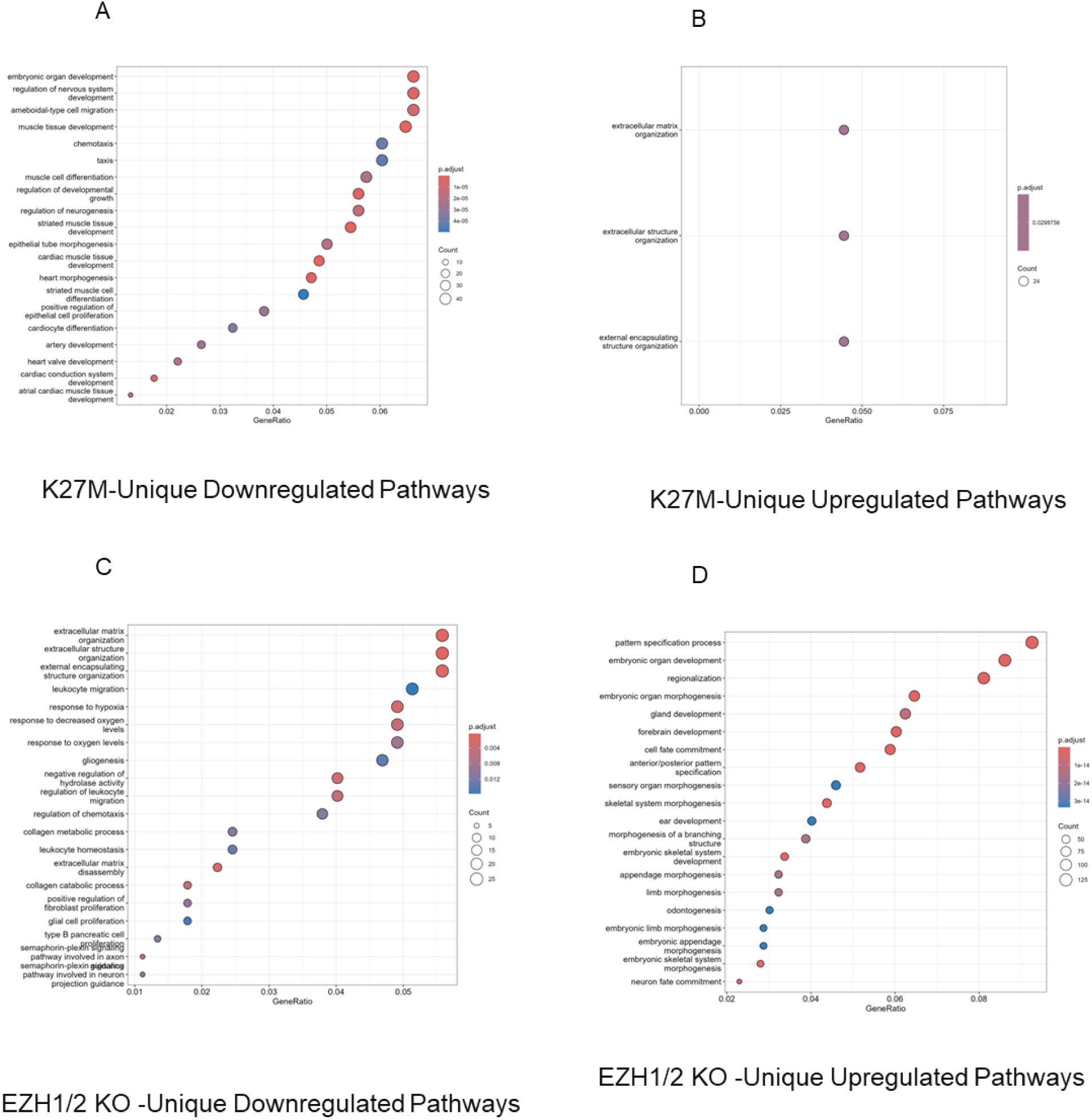
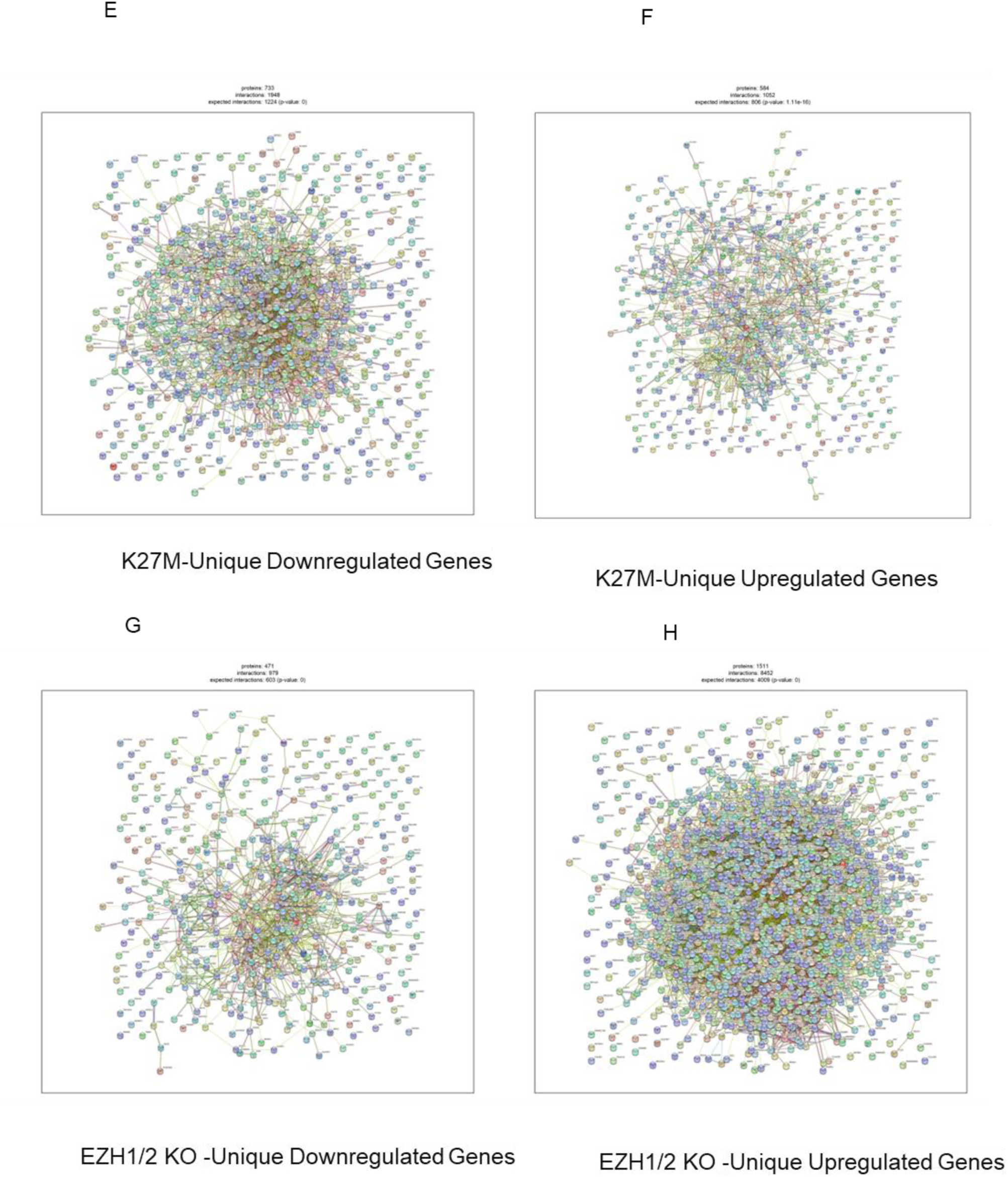
Summary of distinct GO enriched pathways and gene networks associated with K27M versus EZH1/2 KO. The figure presents a Gene Ontology (GO) enrichment analysis dot plot in K27M unique, and EZH1/2 unique genes in SF8628 cells. Each dot represents an enriched GO term, with the dot’s size reflecting the gene count associated with that term and its color indicating the level of statistical significance (p-adjust value). The x-axis, denoting GeneRatio, shows the proportion of genes associated with each GO term relative to the total number of genes analyzed. Only top 20 significantly enriched biological processes are shown. (A) Unique downregulated genes (886 genes) in K27M (B) Unique upregulated genes (813 genes) in K27M (C) Unique downregulated genes (673 genes) in EZH1/2 knockout (D) Unique upregulated genes (1997 genes) in EZH1/2 knockout. (E-H) This panel illustrates the network interactions of genes uniquely regulated in K27M mutants and EZH1/2 knockout SF8628 cells. Nodes represent proteins encoded by differentially expressed genes, while edges indicate protein-protein interactions with a confidence score threshold of 400. Networks are generated using STRING database with a statistical significance annotation based on p-value. E) Downregulated genes in SF8628 K27M cells show a distinct network pattern, pointing to inhibited pathways in the presence of the mutation. F) Upregulated genes in SF8628 K27M cells display a network of enhanced protein interactions, suggesting active signaling pathways specific to the K27M mutation. G) Downregulated genes in SF8628 EZH1/2 knockout cells provide insights into the pathways suppressed due to the loss of EZH1/2 function. H) Upregulated genes in SF8628 EZH1/2 knockout cells reveal a network indicating the molecular effects of PRC2 disruption on cellular signaling.

The difference in the number of enriched pathways—only three for upregulated genes compared to the the 360 for downregulated (top 20 shown)—underscores the mutation’s selective impact on gene expression. The significant upregulation of extracellular matrix-related genes, despite a general trend of repression seen in other developmental pathways, could indicate a dual role of K27M: it not only represses genes critical for controlled cell growth and differentiation but also simultaneously activates specific pathways that may contribute to the malignant phenotype by remodeling the tumor microenvironment.

The GO enrichment analysis and gene network visualization reveal a profound shift in cellular dynamics upon EZH1/2 knockout, reflective of PRC2 loss. The GO dot plot for upregulated genes showcases a predominance of developmental processes, with top pathways such as pattern specification, organ development, and cell fate commitment coming to the fore out of a total of 854 enriched pathways (Figure 5D). This indicates a reawakening of developmental gene programs typically silenced by PRC2 in differentiated cells. Comparatively, the downregulated gene pathways are fewer, with 31 pathways significantly enriched. These include crucial structural and stress response elements like extracellular matrix organization, indicating a downregulation of genes involved in maintaining cellular structure and responding to environmental stressors (Figure 5C).

The complexity of these shifts is visually captured in the network plots. The upregulated genes form a dense network with an unexpected number of interactions (Figure 5H), suggesting a broad and intricate engagement of developmental pathways post-PRC2 knockout. On the contrary, the downregulated genes present a sparser network, implying a lesser degree of interaction and a possible dismantling of cellular architecture and stress response capabilities (Figre 5G).

The loss of PRC2 leads to the activation of developmentally programmed genes while simultaneously reducing the expression of genes responsible for structural maintenance and environmental resilience.

### Summary of stem cell genes, oncogenes and tumor suppressor genes

In our analysis, the CDKN2A gene was significantly upregulated in K27M mutants when compared to their isogenic wild-type controls. Specifically, the SF8628 K27M, SF 8628 K27M EZH1/2KO, and DIPG XVII K27M cell lines showed log2FC of 12.9, 13.4, and 10.9, respectively (Figure 6A). In contrast, the EZH1/2 knockout cells without K27M mutation did not exhibit changes in CDKN2A expression, suggesting a K27M-specific regulatory effect independent of PRC2’s methylation activity.

**Figure 6.**
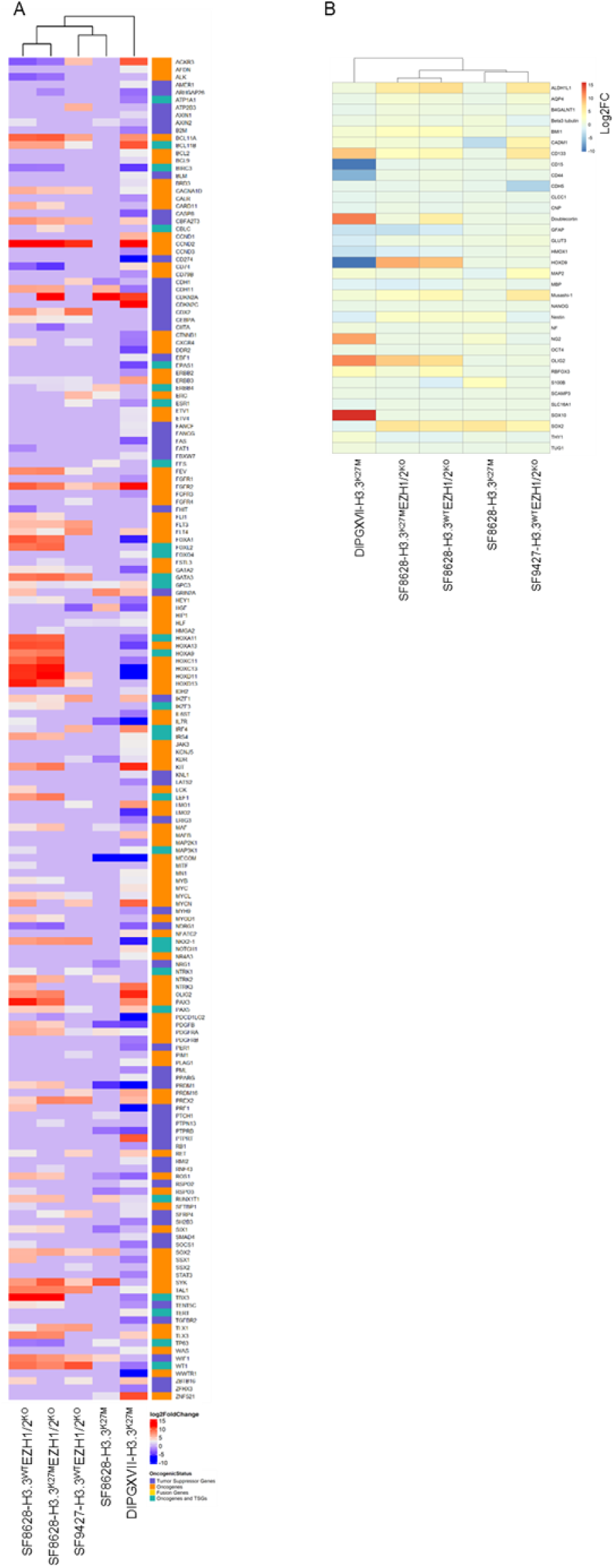
Summary of TSG/Oncogenes/ regulated by K27M. Heatmap displaying differential expression of known oncogenes and tumor suppressor genes from the Cancer Gene Census Tier 1 (as retrieved from Cosmic) in K27M and PRC2 mutant cell lines relative to their isogenic wild-type counterparts. The heatmap illustrates log2 fold change (log2FC) values, with red indicating upregulation and blue indicating downregulation in the mutants. Genes included in this figure were stringently selected for a log2FC greater than 1.5. Cells marked with zero indicate absence of data for a given gene in one of the mutation types, replacing non-available (NA) entries for clearer visual representation. This heatmap highlights the oncogenes and tumor suppressor genes most significantly influenced by the mutations, shedding light on the genetic disruptions driving DMG pathogenesis.

Hepatocyte growth factor (HGF) is known for its role in promoting cell proliferation, migration, and angiogenesis in glioma (39). In our study, we observed a marked variation in HGF expression across different cell lines carrying the K27M mutation or EZH1/2 knockouts. Western blot analysis revealed that HGF expression was notably high in SF8628 K27M mutant cells, significantly reduced in wild-type revertants, and absent in EZH1/2 knockout cells (Supplementary Figure 5B). This pattern indicates that HGF expression is closely associated with the K27M mutation rather than the loss of EZH1/2 or alterations in H3K27me3 levels. Furthermore, treatment with the EZH2 inhibitor EPZ6438 in SF8628 cells led to a partial reduction in HGF expression (Supplementary Figure 5C), suggesting that EZH2 activity is a regulator but not the sole controller of HGF expression. RNA-seq further demonstrated this effect, with a notable increase in HGF expression (log2FC = 5.5) in SF8628 K27M cells versus wild-type counterparts (Figure 6A), while DIPG K27M cells and SF9427 EZH1/2 knockout cells exhibited decreased levels (log2FC = −4.5 and −3.9, respectively).

### Pharmacological EZH2 inhibition

To further validate the effects of EZH1/2 knockout, we examined the impact of the small molecule EZH inhibitor EPZ-6438. This compound selectively inhibited K27 methylation within the 1-10µM concentration range without causing off-target toxicity, as shown in Supplementary Figure 3A,B. MTS assays over 72 hours indicated minimal effect on cell proliferation (Supplementary Figure 3C). Extending the investigation to 21 days, EPZ-6438 significantly reduced proliferation in H3.3K27M mutant cells compared to wild-types (Supplementary Figure 3D). EZH1/2 knockout cells showed resistance to this inhibition (Supplementary Figure 3E), confirming the specificity of our gene editing and EPZ-6438’s selectivity.

Western blots assessed neural stem cell protein expression in EZH1/2 knockout and control cells, and cells treated with 10µM EPZ-6438 for 72 hours. Analyzed proteins included SOX2, Olig2, Nestin, Nanog, GFAP, H3K27M, H3K27Me3, histone H3, and tubulin (Supplementary Figure 5A). Significant changes in protein profiles suggested EZH2 enzymatic influence on neural stem cell networks, with notable adaptations in Olig2, Nestin, Nanog, and GFAP occurring after EZH1/2 deletion but not after pharmacological inhibition, indicating longer-term effects.

RNA sequencing and Western blot analyses were conducted to investigate HGF expression dysregulation in H3.3K27M mutants and controls, including cells treated with 10µM EPZ-6438 for 72 hours and untreated EZH1/2 knockout and control cells (Supplementary Figure 5B,C). These studies confirmed HGF regulation by the K27M mutation, independent of H3 K27me3 loss.

### In Vivo Evaluation of Tumor Growth and Progression in Xenograft Model

To evaluate the tumor-initiating capabilities of genetically modified cell lines, immunodeficient nude mice were inoculated with 1 × 10^6 cells from various H3.3 and EZH1/2 CRISPR-modified lines including SF8628 H3.3K27M, SF8628 H3.3 WT (pooled clones), SF8628 H3.3K27M EZH1/2 knockout (pooled clones), SF8628 H3.3 WT knockout (pooled clones), SF9247 H3.3 WT, and SF9247 EZH1/2 knockout clones. Among these, only SF8628 H3.3K27M cells initiated tumor growth (Supplementary Figure 6), with other lines failing to form tumors. At 200 days, insufficient tumor-bearing mice reached the experimental endpoint of 2000 mm3 to declare a significant difference in overall survival.

Further studies explored the effects of CRISPR-induced EZH1/2 deletions on tumor growth using adult glioblastoma multiforme (GBM) xenograft models, U87 and U118. Both EZH1/2 knockout (3 pooled clones) and control variants of these lines were tested, showing that EZH1/2 knockout significantly delayed tumor onset and increased survival, without changing the growth rate of the tumors in vivo (Supplementary Figure 7,8).

## Discussion

The mechanism by which H3K27M exerts its derepressing effect on gene regulation and promotes tumorigenesis remains a topic of debate. It is thought that the derepression of genes through the global loss of K27me3 in K27M cells is responsible for the associated oncogenic transformation. Following that line of thinking, many competing molecular models have been proposed (each with supporting experimental evidence) to explain how the K27M mutation causes the depletion of K27me3. These theories propose that H3.3K27M mutation involves either the sequestration of PRC2 (24–26,40), selective retention of PRC2 activity at specific sites (29), exclusion of PRC2 from certain chromatin regions (30), an allosteric modification (poison model) that impacts PRC2’s distribution and function across the genome (31), or a ‘step down’ gradation from me3 to me2 and me1 (32), suggesting complex interactions that disrupt traditional methylation patterns. Our data, however, adds a new level of complexity to these models as it suggests a unique epigenetic function to K27M, itself, that is necessary for the tumor formation and likely works synergistically with K27me3 depletion in glial cell transformation.

Research on small molecule inhibitors of EZH2 has demonstrated potential for reducing H3K27me3 (29,30,41,42), paradoxically leading to decreased methylation at silenced tumor-suppressor genes in DMG and inhibiting cell proliferation (29,30). Our findings corroborate this approach, as both PRC2 knockout mice and K27M revertants showed no tumor development, highlighting the critical role of precise methylation balance “just right model” in tumorigenesis. Deviations from this balance reduce tumorigenic potential, offering therapeutic opportunities. In our study, PRC2 knockout in SF8628 cells prevented tumor formation in mice, in contrast to SF8628 K27M cells, which formed tumors. This supports Mohammad et al.’s findings (29), on the essential role of PRC2 in tumor proliferation in H3K27M-expressing tumors.

Lewis et al., 2022, (38) report that the H3.3K27M mutation in DIPG lines SU-DIPG-XIII and SU-DIPG-XVII, and their gene-edited wild-type counterparts, leads to open chromatin that enhances expression of crucial developmental genes like ASCL1 and NEUROD1, potentially accelerating tumor progression. Their findings emphasize the mutation’s impact on chromatin dynamics and gene regulation, notably enriching transcription factors such as ASCL1, NEUROD1, OLIG2, and HOXA2 in K27M mutants. In contrast, our study noted a significant upregulation of ASCL1 in K27M DIPG 17 cells (log2FC = 11.8) without a parallel increase in SF8628 K27M or SF9427 PR2KO cells. NEUROD1, OLIG2, and HOXA2 also showed increased levels in K27M DIPG 17 but not SF8628 cells. These observations suggest that the K27M mutation’s effects on gene expression and chromatin dynamics are cell-type specific and heavily influenced by the cellular context (Fig 7b). In our study, we observed significant upregulation of the CDKN2A gene in K27M mutant models such as SF8628 K27M, SF8628 K27M EZH1/2 KO, and DIPG XVII K27M, with respective log2 fold changes of 12.9, 13.4, and 10.9. This upregulation was notably absent in EZH1/2 knockout models lacking the K27M mutation, suggesting that K27M exerts its regulatory effects on CDKN2A independently of PRC2 activity. In contrast, our ATAC-seq profiles show that HOX genes, typically regulated by PRC2, remain comparatively inaccessible in K27M cells, and become accessible only upon PRC2 knockout, indicating a strong regulatory control by PRC2 rather than histone modification changes induced by H3K27M.

Chen et al., (44) observed that the K27M mutation generally decreases H3K27me3 levels, predominantly leading to gene induction with 139 genes upregulated across K27M lines. In contrast, our study finds that K27M not only interacts with PRC2 but also affects gene regulation more evenly, both upregulating and downregulating genes, with a more significant impact on the downregulation of biological pathways, imposing an overall repressive effect on the transcriptome. This observation indicates that previous studies comparing K27M effects solely with wild-type controls may have missed the mutation’s complex role due to overlapping PRC2 impacts on gene regulation. Our results reveal that K27M alters PRC2 function and independently influences gene expression, suggesting a dual mechanism where K27M can both modify and counteract PRC2 activity, leading to varied regulatory effects on the transcriptome.

We observed that the reversion to wildtype in H3.3K27M mutant cells led to significant reductions in growth and survival over 21 days. The well-documented response of K27M mutant cells to EZH2 inhibition was confirmed in vivo; however, this response was reversed in K27M wildtype revertants, and EZH2 knockouts were found to be insensitive to EZH1/2 inhibition with EZP. Notably, both K27M wildtype revertants and EZH1/2 knockout cells failed to grow in immunodeficient mice, unlike K27M mutant cells. Furthermore, two well-studied glioblastoma cell lines showed a very significant reduction or delay in growth in immunodeficient mice following EZH2 deletion.

The differential responses observed between H3.3K27M, H3.3 wildtype revertants, and EZH2 knockout cells emphasizes the potential specificity and intimal effectiveness of EZH2- targeted therapies in K27M-mutated scenarios. Moreover, the inability of H3.3 WT revertant and EZH1/2 knockout cell lines to proliferate in immunodeficient mice suggests that the K27M mutation may confer unique growth advantages independent from the loss of H3K27me3.

In summary, our findings highlight the K27M mutation’s multifaceted impact on gene expression, chromatin accessibility, and tumor dynamics, which transcends mere reductions in H3K27me3. The mutation introduces a dual regulatory mechanism—modulating and counteracting PRC2—that significantly influences gene expression and biological pathways. Notably, this is the first evidence of genes uniquely deregulated by K27M, independent of K27 methylation loss, show marked changes in chromatin accessibility. This specific pattern, distinct from changes driven by EZH1/2 knockouts, defines H3.3K27M’s unique effect on chromosomal accessibility, independent of its impact on chromatin via loss of K27 methylation.

## Author contributions

SB analyzed data and wrote the manuscript. FLH and CAD performed the study. FG assisted with data analysis and study design. AL and IE supported the research. EHH supervised CAD and AL and provided reagents. JPR, as Principal Investigator, designed the study, supervised the research, and wrote the manuscript.

## Data Availability

The data supporting the findings of this study are available from the corresponding author upon reasonable request.

## Acknowledgment

We thank Paul Knoepfler (UC Davis School of Medicine, Sacramento, CA) for kindly providing the DIPGXVII DMG H3.3 K27M mutant and wild-type revertant cells.

