## Supplementary material for "H3F3A K27M Mutations Drives a Repressive Transcriptome by Modulating Chromatin Accessibility, Independent of H3K27me3 in Diffuse Midline Glioma": supp tables and figures

**Supplementary Table 1. Mass spec analysis of histone PTM at K27, in DMG with histone mutations**. There are two copies of H3.3, three copies of H3.2, and 10 copies of H3.1 genes (41). The relative abundance of each form of histone PTMs. The presence of the K27M mutation was also detected and was effectively removed by CRISPR-Cas9 in the revertant cell line.

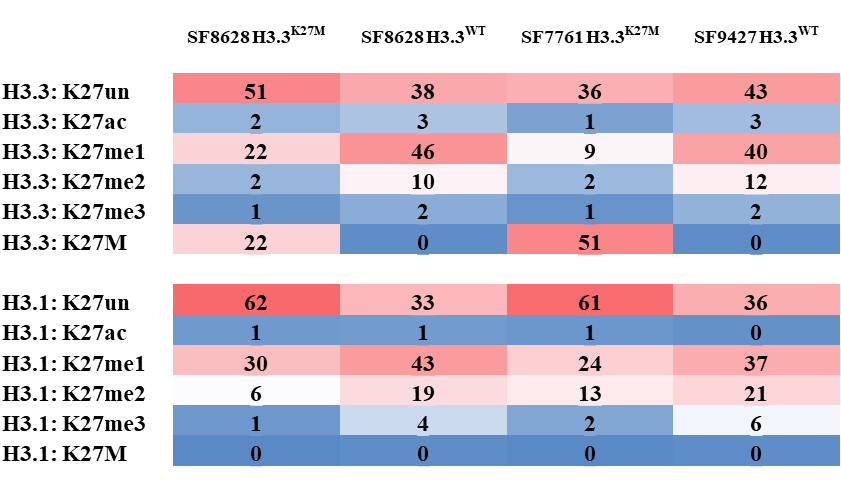

Displayed values indicate the percentage of each histone modification relative to the total pool of the corresponding tryptic peptide across all samples. "Un" denotes unmodified histones; "ac" indicates acetylation; "me1," "me2," and "me3" refer to mono-, di-, and tri-methylation, respectively. Data points are the average of three mass spectrometry runs, with standard deviations uniformly below 0.1% and thus not displayed in the table. Colors in the table highlight relative abundances: red signifies higher and blue lower relative abundances compared to other samples.

**Supplementary Table 2: Summary of the number of genes and pathways dysregulated by K27M and EZH1/2 knockout.**

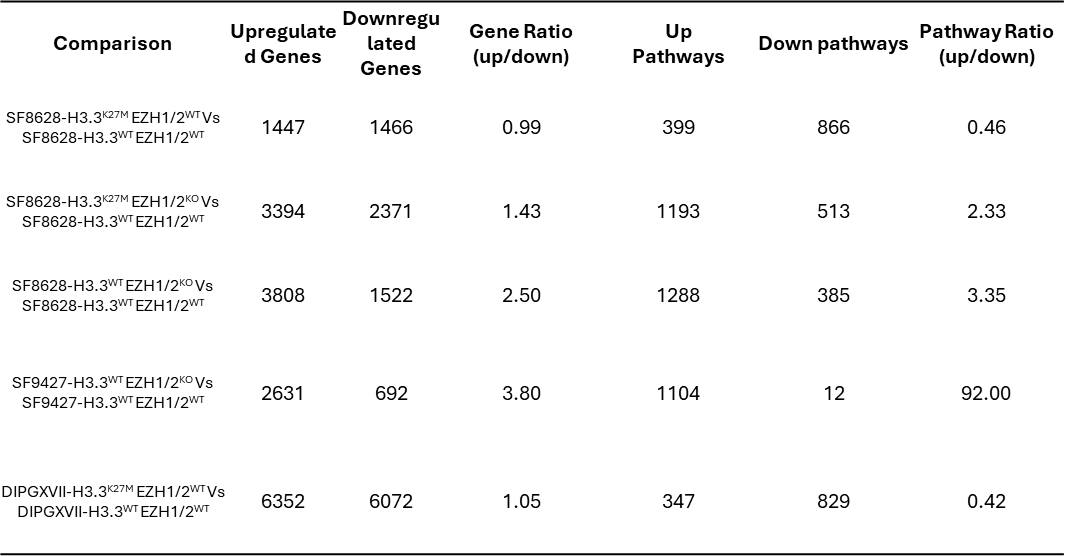

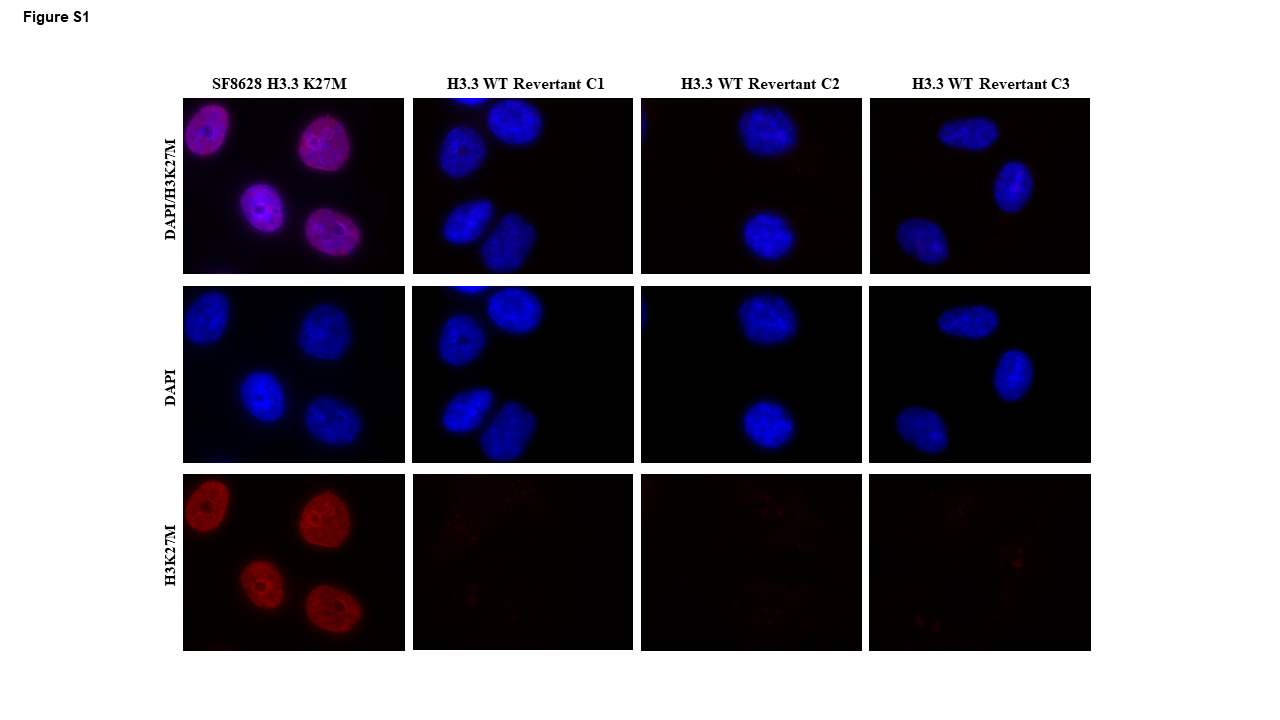

**Supplementary Figure 1: Immunofluorescence panel; confirmation of H3.3K27M revertants**. This panel illustrates immunofluorescence staining demonstrating loss of H3.3K27M at histone H3 lysine 27 (H3K27me3) in SF8628 CRSIPR revertant clones compared to wildtype cells. Red fluorescence indicates H3.3K27M staining, while blue represents nuclear counterstaining with DAPI.

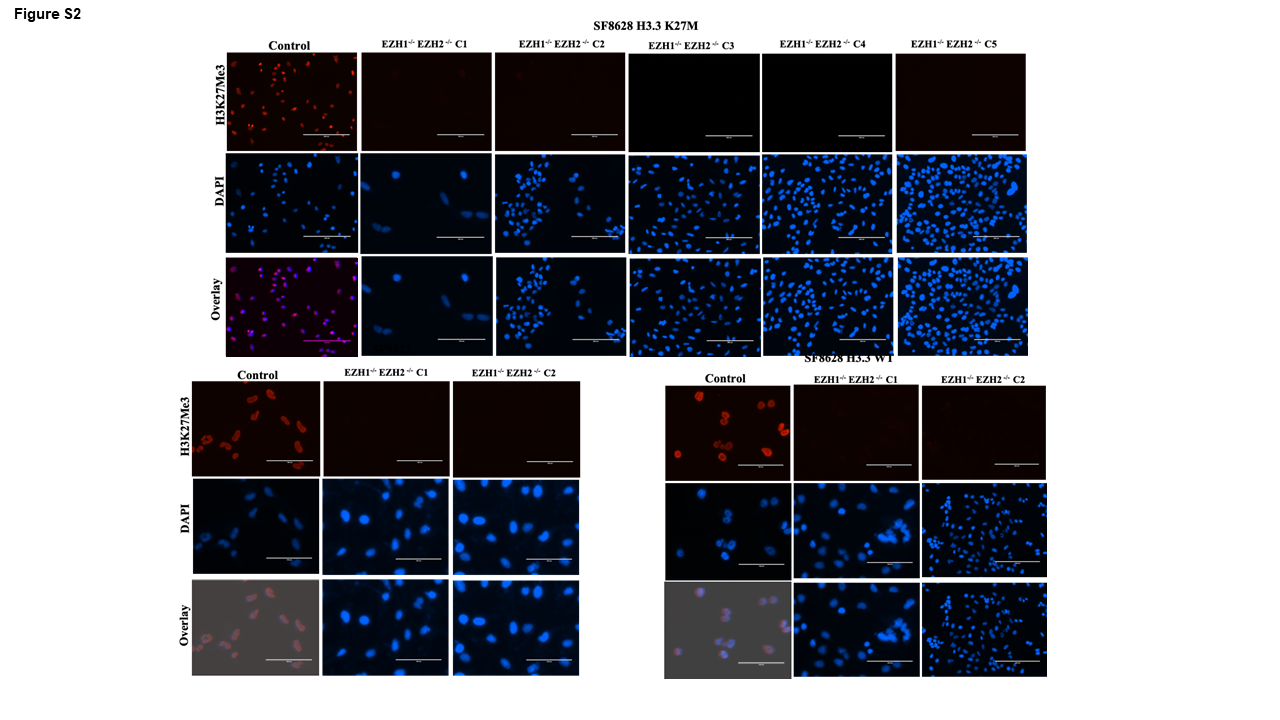

**Supplementary Figure 2: Immunofluorescence panel; confirmation of EZH1/2 knockout.** This panel illustrates immunofluorescence staining demonstrating the reduction of trimethylation at histone H3 lysine 27 (H3K27me3) in EZH1 and EZH2 cells compared to wildtype cells. Red fluorescence indicates H3K27me3 staining, while blue represents nuclear counterstaining with DAPI.

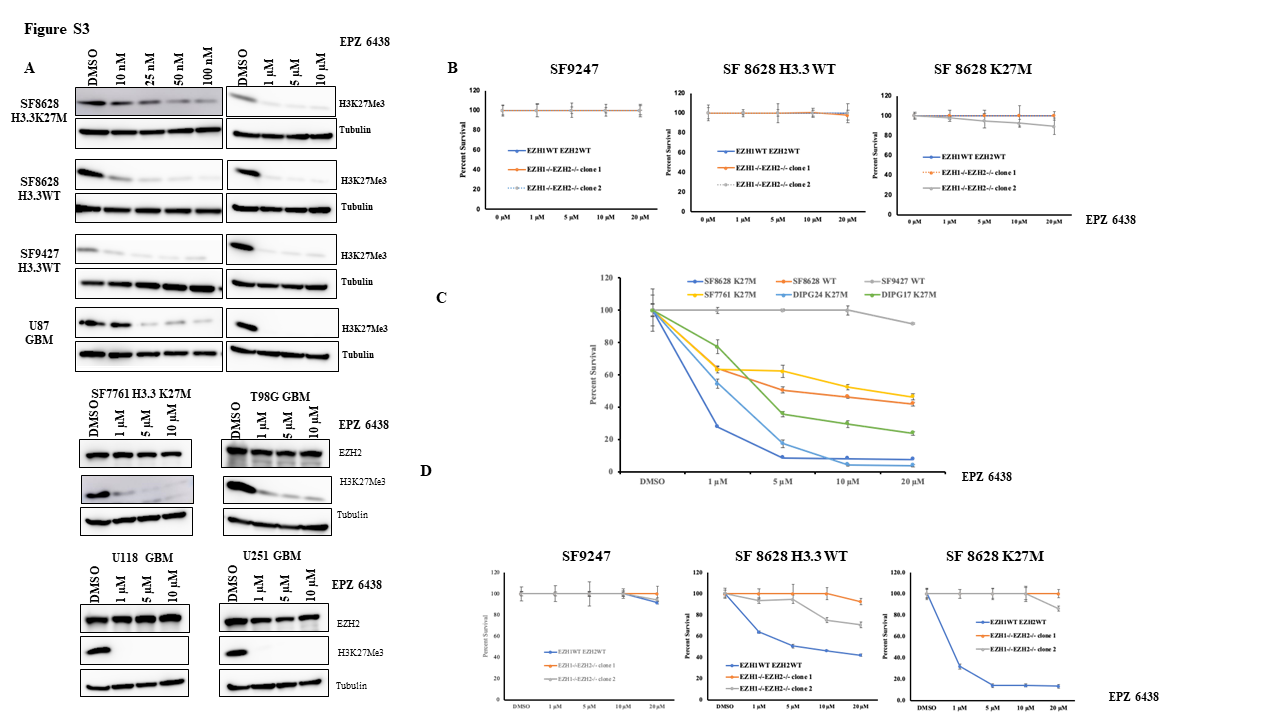

**Supplementary Figure 3: Effects of Tazemetostat (EPZ-6438) on H3.3K27M Mutant, Revertant, and EZH1/2 Knockout Cells** (a,b) Western Blot Analysis: Dose-dependent impact of EPZ6438 (10 nM to 10 µM) on H3.3 lysine 27 trimethylation (H3.3K27Me3) levels in cells with wild-type EZH1/2.

(c) 72-Hour MTS Assay: Viability of parental and EZH1/2 knockout cells following treatment with EPZ-6438 across various concentrations, assessed over a 72-hour period. (d) Long-Term Cell Survival Assay (21 Days): Cells were treated with EPZ-6438 for two weeks. Post-treatment, colonies were stained with crystal violet, and absorbance at 450 nm was measured to evaluate cell survival**.** (e) Response of EZH1/2 Knockout Cells to Treatment: This panel demonstrates that the loss of EZH1/2 enhances the efficacy of EPZ-6438, inhibiting cell growth through simultaneous inhibition of EZH1 and EZH2 pathways.

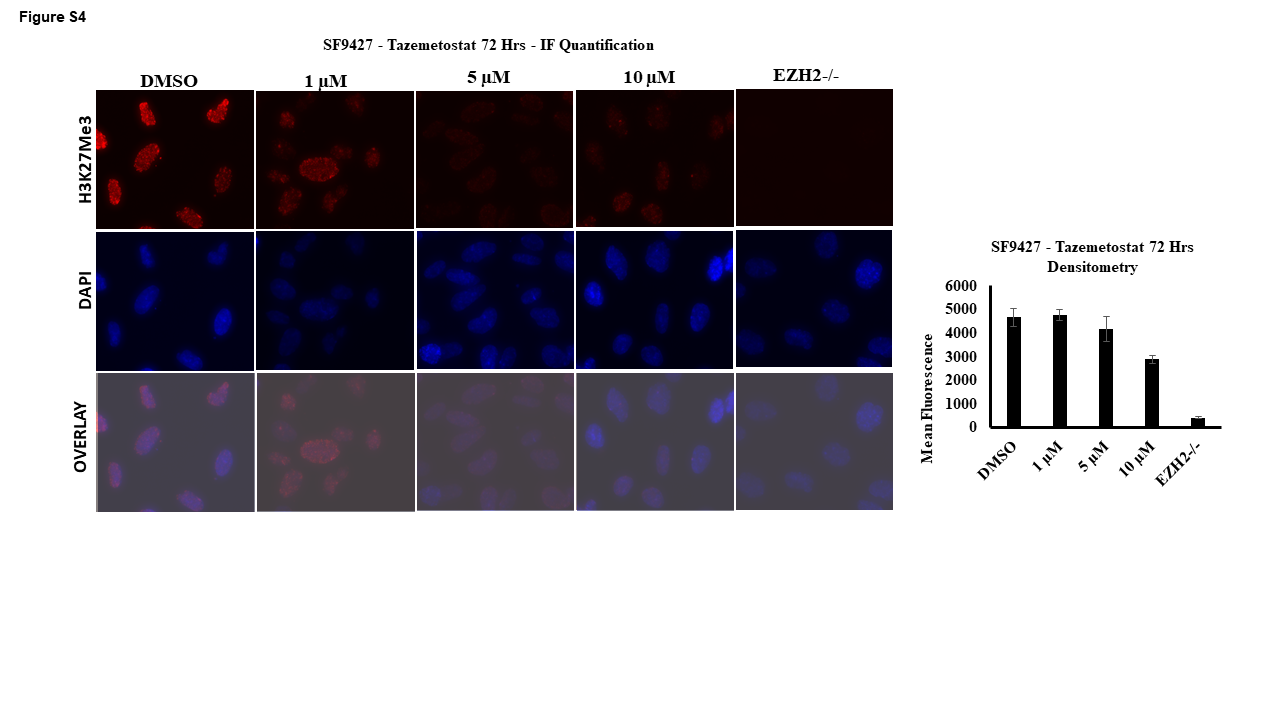

**Supplementary Figure 4: Dose-dependent Effects of Tazemetostat (EPZ6438) on H3K27 Trimethylation in SF9247 Cells**

This panel displays immunofluorescence staining that quantifies the reduction of trimethylation at histone H3 lysine 27 (H3K27me3) in response to treatment with EPZ6438, ranging from 1 µM to 10 µM. The reduction is compared to EZH1/2 knockout control cells. Red fluorescence indicates H3K27me3 staining, and nuclei are counterstained in blue with DAPI.

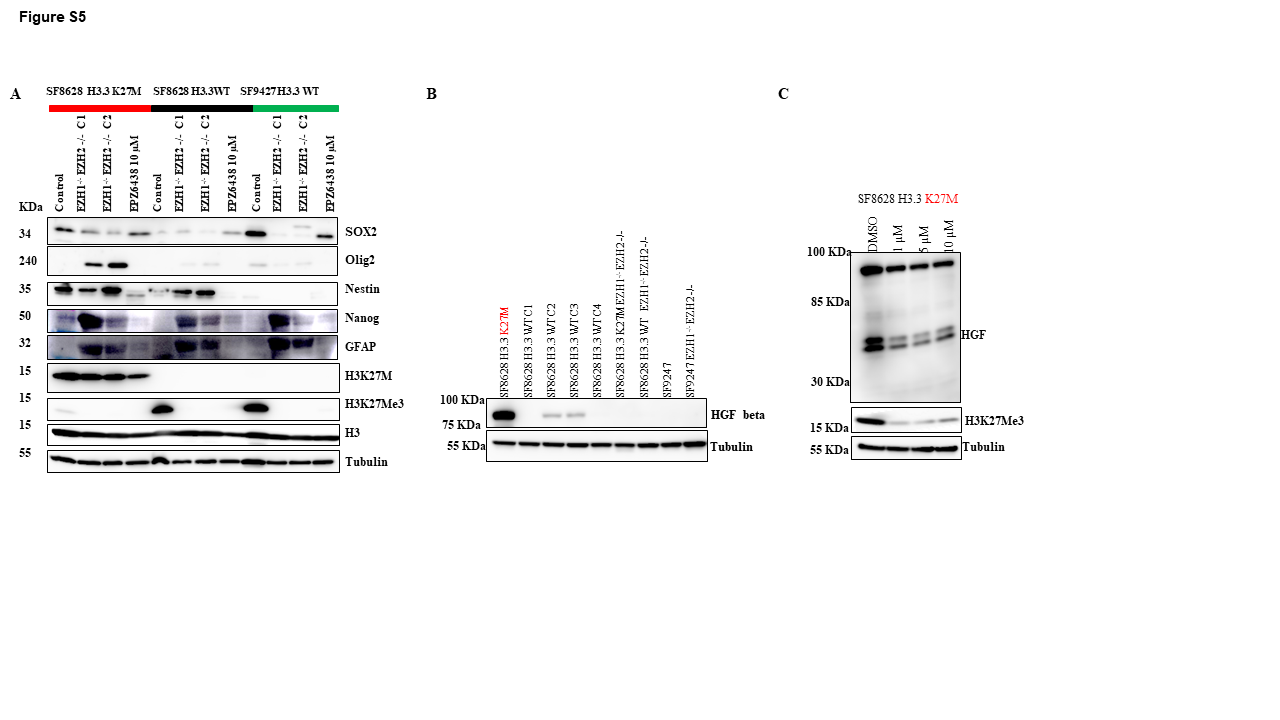

**Supplementary Figure 5: Effects of H3.3K27M Reversion, EZH1/2 Knockout, and EZH2 Inhibition on Stem Cell Marker Expression in DMG Cells**

(a) Western Blot Analysis: This panel illustrates the impact of CRISPR-Cas9 homology-directed repair (HDR)-mediated reversion of the H3.3K27M mutation to wild type and knockout of EZH1/2, which reduces H3K27 methylation. The blot shows changes in the expression levels of various stem cell markers and histone modifications in response to genetic modifications and EZH2 inhibition:

SOX2: Marker of pluripotency and neural stem cells.

Olig2: Associated with neural progenitor cells.

Nestin: Intermediate filament protein expressed in stem cells.

Nanog: Essential for the self-renewal of undifferentiated embryonic stem cells.

GFAP: Marker for astrocytes in the central nervous system.

H3K27M: Mutant histone protein.

H3K27Me3: Marker of transcriptional repression.

H3 and Tubulin: Used as loading controls.

(b) HGF Expression Analysis in SF8628 Cells: This panel links HGF expression primarily to mutant H3.3K27M rather than to the loss of EZH1/2 or changes in H3K27Me3 levels.

(c) EPZ6438 Treatment Impact on HGF Expression: Despite treatment with EPZ6438, complete repression of HGF expression in SF8628 cells was not achieved.

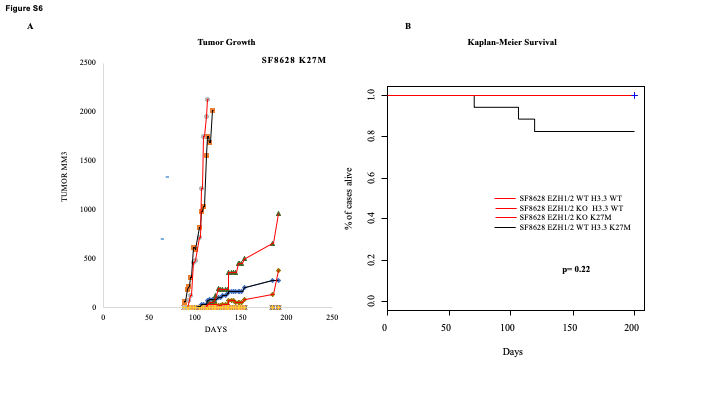

**Supplementary Figure 6: Impact of EZH1/2 Knockout on DMG Cell Growth In Vivo.** To assess the tumor-initiating capabilities of genetically modified cell lines, immunodeficient nude mice were injected with 1 × 10^6 cells from various H3.3 and EZH1/2 CRISPR-modified lines. SF 8628 H3.3 K27M, SF 8628 H3.3 WT (pooled clones), SF 8628 H3.3 K27M EZH1/2 knockout (pooled clones), SF 8628 H3.3 WT knockout (pooled clones), SF 9247 H3.3 WT, and SF 9247 EZH1/2 knockout clones. (a)Of these, only the SF 8628 H3.3 K27M cells-initiated tumor growth, with a mean latency of 180 days. The other cell lines did not result in tumor formation. (b) Kaplan-Meier Survival Estimate: Provides a survival analysis showing that EZH1/2 knockout in the indicated cells lines did significantly increases survival (p<0.022) due to the majority tumor bearing mice still being alive at the experimental censor date.

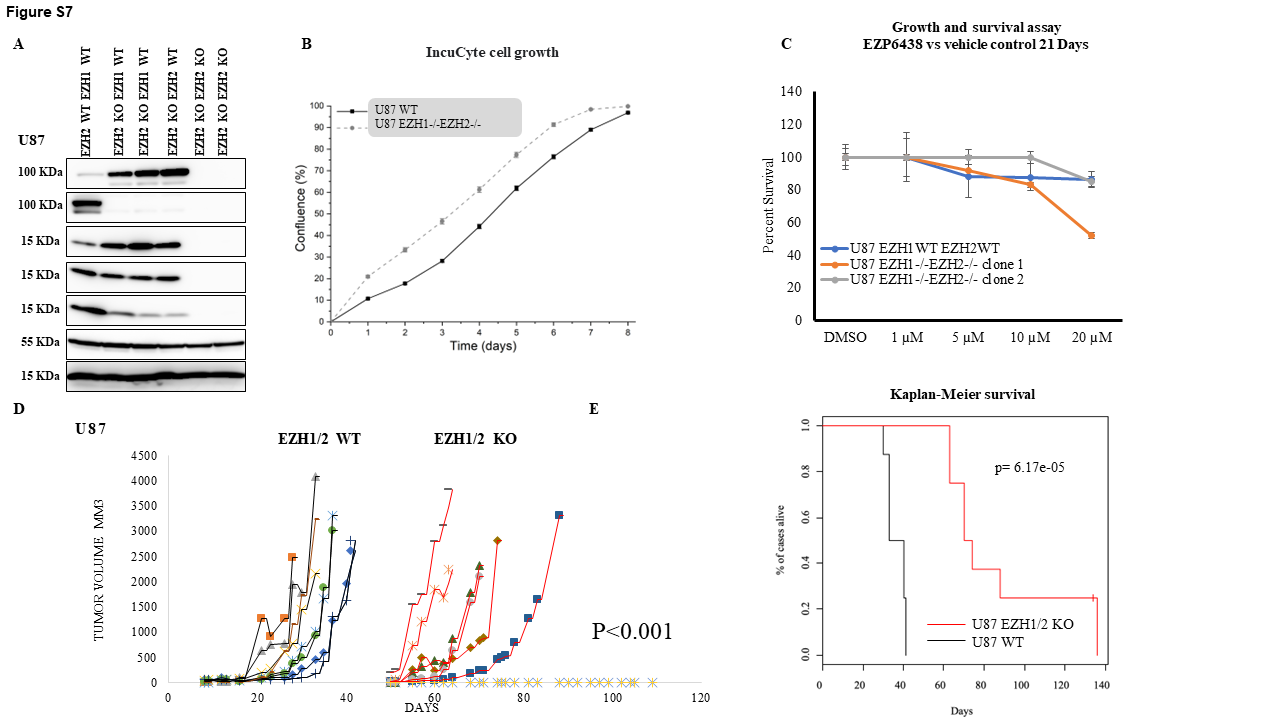

**Supplementary Figure 7: Impact of EZH1/2 Knockout on In Vivo and In Vitro Growth of U87 Glioma Cells.** (a) Western Blot Analysis**:** This panel shows the decrease in histone H3 lysine 27 (H3K27) methylation following CRISPR-Cas9-mediated knockout of EZH1 and EZH2 in U87 glioma cells, confirming effective enzymatic inhibition. (b) Growth Rate Analysis of H3.3 Wildtype Revertant Clones: This section evaluates the proliferation rates of U118 cells that have reverted to wildtype H3.3 status, with comparisons to unmodified control cells. (c) Cell Growth and Viability Over 21 Days: Assesses the long-term viability and growth rates of parental and EZH1/2 knockout U87 cells under standard culture conditions.(d) In Vivo Tumor Growth: Details the experimental setup where two clones of U118 cells, with and without EZH1/2 knockout, were injected subcutaneously into the flanks of cohorts of eight immunodeficient nude mice. It reports a significant reduction in tumor growth in mice injected with EZH1/2 knockout cells (P < 0.001).

(e) Kaplan-Meier Survival Estimate: Provides a survival analysis showing that EZH1/2 knockout significantly increases survival (p<0.001), highlighting the potential therapeutic benefits of EZH1/2 inhibition

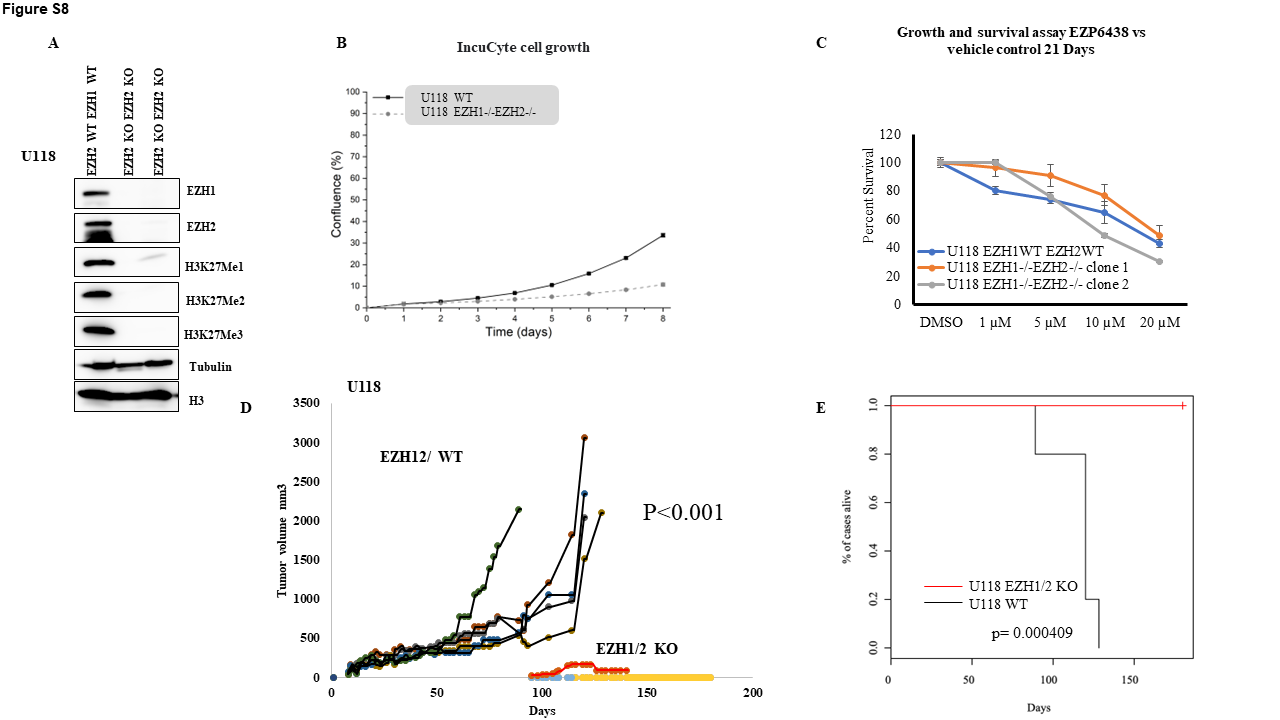

**Supplementary Figure 8: Impact of EZH1/2 Knockout on In Vivo and In Vitro Growth of U118 Glioma Cells.** (a) Western Blot Analysis: This panel shows the decrease in histone H3 lysine 27 (H3K27) methylation following CRISPR-Cas9-mediated knockout of EZH1 and EZH2 in U118 glioma cells, confirming effective enzymatic inhibition. (b) Growth Rate Analysis of H3.3 Wildtype Revertant Clones: This section evaluates the proliferation rates of U118 cells that have reverted to wildtype H3.3 status, with comparisons to unmodified control cells. (c) Cell Growth and Viability Over 21 Days: Assesses the long-term viability and growth rates of parental and EZH1/2 knockout U118 cells under standard culture conditions. (d) In Vivo Tumor Growth: Details the experimental setup where two clones of U118 cells, with and without EZH1/2 knockout, were injected subcutaneously into the flanks of cohorts of eight immunodeficient nude mice. It reports a significant reduction in tumor growth in mice injected with EZH1/2 knockout cells (P < 0.001). (e) Kaplan-Meier Survival Estimate: Provides a survival analysis showing that EZH1/2 knockout significantly increases survival (p<0.001), highlighting the potential therapeutic benefits of EZH1/2 inhibition

**Supplementary methods**

**Western blotting:** Primary antibodies were obtained from the following sources and used according to manufacturer’s recommendations; Anti-EZH1 (42088, 1:1,000, Cell Signaling Technology, Danvers, MA), Anti-EZH2 (5246, 1:1000, Cell Signaling Technology), Anti-Mono-Methyl-Histone H3 Lys27 antibody (84932, 1:1000, Cell Signaling Technology), Anti-Di-Methyl-Histone H3 Lys27 antibody (9728, 1:1000, Cell Signaling Technology), Anti-Tri-Methyl-Histone H3 Lys27 antibody (9733, 1:1000, Cell signaling Technology), Anti-Histone H3 mutated K27M antibody (ab190631, 1:1000, Abcam), Anti–Histone 3 (9715, 1:1000, Cell Signaling Technology), and anti-β-tubulin (ab21058, 1:5000, Abcam). The blots were then incubated with an anti-mouse or anti-rabbit IgG-HRP secondary antibody (7074, 7076 Cell Signaling Technology), incubated with ECL solution (Amersham, GE Healthcare, Chicago, IL), and imaged on a AI600 Chemiluminescent Imager (Amersham). (DAPI) diluted in TBS for 5 min. Coverslips were mounted using ProLong Gold anti-fade (Life Technologies, Carlsbad, CA).

**Mass spec Quantification of Histone modifications**

Histones were acid extracted, derivatized via propionylation, digested with trypsin, newly formed N-termini were propionylated as previously described (59), and then measured 3 seperate times using the Thermo Scientific TSQ Quantum Ultra mass spectrometer coupled with an UltiMate 3000 Dionex nano-liquid chromatography system. The data was quantified using Skyline (60), and represents the percent of each modification within the total pool of that tryptic peptide.

**Growth Curves.** Cells were plated into 24 well dishes in DMEM with 10% FBS. Growth was monitored on an IncuCyte S3 in incubator microscope at 37°C and 5% CO2. Cells were imaged in white light every 4 hours using a 10x lens. IncuCyte software was used to calculate percentage confluency and data exported in text form. Growth curves were then regraphed in Origin 2018b (OriginLab).

**Immunofluorescence**. Primary antibodies anti-histone H3 K27M, (31-1175-00, 1:10,000, RevMAb Bioscience), and anti-histone H3 K27me3 (9733, 1:10,000, Cell signaling Technology), Cells were washed three times for 10 minutes each and then indirect fluorescence was achieved using Alexa 488, 594, or 660 conjugated goat secondary antibodies (Life Technologies) for 3 hours at room temperature. Cells were washed three times for 10 minutes with 1% BSA/0.05% Tween TBS and incubated with 4′, 6-diamidino-2-phenylindole

**Xenograft assays**. Parental and EZH1/2 knockout cells of SF8628, B23, SF9427, U87 and U118 were harvested and resuspended in cold PBS and Matrigel (1:1 ratio). One million cells were injected using a 1 mL syringe (BD Biosciences, USA) with 22G x 1-1/2 needle (Covidien, USA) in each flank of NU/J mice (Jackson Labs). Mice were visually monitored for tumor formation and measured using vernier caliper. All mouse protocols were approved by, University of Minnesota Institutional Animal Care and Use Committees.

**MTS assay.** 10,000 cells were seeded in 96 well plate and allowed to adhere over-night. Cells were treated with either DMSO or EZH2 inhibitor EPZ6438 at different concentrations (1, 5, 10, 20 µM) for 72 hrs. Cell viability measured using MTS reagent (Abcam, USA).

**Clonogenic assay**. 3,000 cells were seeded in 6 well plates and allowed to adhere over-night. Cells were treated with DMSO or EZH2 inhibitor EPZ6438 for 21 days. Then cells were fixed and stained with crystal violet containing methanol. Crystal violet dissolved in 10% acetic acid and absorbance measured at 450 nM.
